# Barcoded monoclonal embryoids are a potential solution to confounding bottlenecks in mosaic organoid screens

**DOI:** 10.1101/2025.05.23.655669

**Authors:** Samuel G. Regalado, Chengxiang Qiu, Jean-Benoît Lalanne, Beth K. Martin, Madeleine Duran, Cole Trapnell, Aidan Keith, Silvia Domcke, Jay Shendure

## Abstract

Genetic screens in organoids hold tremendous promise for accelerating discoveries at the intersection of genomics and developmental biology. Embryoid bodies (EBs) are self-organizing multicellular structures that recapitulate aspects of early mammalian embryogenesis. We set out to perform a CRISPR screen perturbing all transcription factors (TFs) in murine EBs. Specifically, a library of TF-targeting guide RNAs (gRNAs) was used to generate mouse embryonic stem cells (mESCs) bearing single TF knockouts. Aggregates of these mESCs were induced to form mouse EBs, such that each resulting EB was ’mosaic’ with respect to the TF perturbations represented among its constituent cells. Upon performing single cell RNA-seq (scRNA-seq) on cells derived from mosaic EBs, we found many TF perturbations exhibiting large and seemingly significant effects on the likelihood that individual cells would adopt certain fates, suggesting roles for these TFs in lineage specification. However, to our surprise, these results were not reproducible across biological replicates. Upon further investigation, we discovered cellular bottlenecks during EB differentiation that dramatically reduce clonal complexity, curtailing statistical power and confounding interpretation of mosaic screens. Towards addressing this challenge, we developed a scalable protocol in which each individual EB is monoclonally derived from a single mESC and genetically barcoded. In a proof-of-concept experiment, we show how these monoclonal EBs enable us to better quantify the consequences of TF perturbations as well as ’inter-individual’ heterogeneity across EBs harboring the same genetic perturbation. Looking forward, monoclonal EBs and EB-derived organoids may be powerful tools not only for genetic screens, but also for modeling Mendelian disorders, as their underlying genetic lesions are overwhelmingly constitutional (*i.e.* present in all somatic cells), yet give rise to phenotypes with incomplete penetrance and variable expressivity.

## Introduction

Transcription factors (TFs) play a key role in cell type specification in mammalian systems, including at the earliest stages of development, such as the appearance of germ layers, as well as at later stages, such as the emergence and diversification of organ progenitors^1,2^. Although much has been learned about developmental TFs by knocking them out in mouse models, this strategy is challenged by the inaccessibility of early *in vivo* development. Put another way, the gross phenotypic consequences of an *in vivo* TF knockout (*e.g.* “early embryonic lethality”) are much easier to determine than the succession of molecular and cellular events that give rise to that phenotype. As we consider the goal of systematically assessing <u>all</u> mammalian TFs for their role in cell type specification, it is natural to turn to *in vitro* models of mammalian development, which are advancing quickly in sophistication^3^.

Organoid models, widely used to model the development or homeostasis of mammalian organs, are complex *in vitro* systems that recapitulate many aspects of *in vivo* tissue organization and physiology^4^. Although a few models begin with single cells, most organoid protocols begin by aggregating hundreds to thousands of unrelated precursor cells, *e.g.* embryonic, induced pluripotent, adult, or cancer stem cells^5,6^. The potential of the resulting aggregates to differentiate into complex 3D structures was first demonstrated in the early 1980’s through the derivation of cyst-like structures composed of all three germ layers termed ’embryoid bodies’ (EBs) from mouse embryonic stem cells (mESCs)^7^. In the ensuing decades, it was shown that subjecting stem cell aggregates to defined media conditions can direct their differentiation towards various structures that model specific aspects of *in vivo* biology, *e.g.* brain, heart, retinal or liver organoids^8–11^. More recently, various groups have sought to apply CRISPR and single cell profiling to organoids in order to study the consequences of genetic perturbations in tissue-like contexts^12–15^.

Since stem cell-derived EBs give rise to various cell types from all three germ layers over the course of just a few weeks, they are an attractive *in vitro* model for studying cell fate specification in early mammalian development. Here we set out to leverage EBs to probe the consequences of perturbing each mammalian TF on the specification of germ layers and early cell types. Specifically, we performed experiments in which a single TF was perturbed by CRISPR/Cas9 in each of many mESCs, which were then aggregated, differentiated to EBs, and profiled by scRNA-seq. This screen was ’mosaic’, in that each aggregate/EB derived from multiple cells that were heterogeneous with respect to their encoded TF perturbation. In theory, depending on the specific TF perturbation, this should lead to impeded (or enhanced) development of certain cell fates relative to counterparts bearing other perturbations or non-targeting control guides.

However, although we identified numerous seemingly significant associations between specific TF perturbations and the enhancement or depletion of specific cell types, our results were irreproducible across independent biological replicates. Further investigation revealed that stochastic cellular bottlenecks during EB differentiation confounded our experimental design.

To address this challenge, we developed a novel protocol for the efficient production of <u>monoclonally derived</u>, individually barcoded EBs. With this approach, any barcode or genetic perturbation present in that founding mESC will be present in every cell of the aggregate, EB or organoid that derives from that founder, facilitating the statistical analysis of effects across many ‘individuals’. We then applied this protocol to conduct a more limited TF perturbation screen. Our results highlight the potential of monoclonal, barcoded EBs as a model system for large-scale genetic screens of early developmental processes, as well as for organoid screens more generally.

## Results

### Characterizing mouse EBs as a model system for studying cell fate specification

EB formation, followed by histological evaluation of sectioned EBs, has been used for decades to test the pluripotency of stem cells^16^. Today, single cell molecular profiling technologies allow us to quantify the expression of thousands of genes instead of a few selected markers, thus enabling a more detailed cataloging of cell type composition in these simple organoids. Whereas human EBs have been profiled at single cell resolution in some depth^17^, scRNA-seq studies of mouse EBs to date have profiled comparatively few cells (200-4000) from only early timepoints (4-14 days)^18–22^.

In order to identify the timepoint in EB differentiation with a level of cell type complexity suitable for a TF perturbation screen, we applied scRNA-seq to profile the transcriptomes of 25,937 cells from days 7, 14 and 21 of mouse EB differentiation. Cell types were annotated based on marker gene expression (**Fig. 1a**; **Supplementary Table 1**). A wide range of cell types were observed, with representation of all three germ layers. We found that the epiblast-like population decreases substantially between day 7 and 14 (*e.g.* epiblast & primitive streak: 50.5 → 13.0%), and is replaced by more differentiated endodermal (*e.g.* parietal endoderm: 0.4 → 7.1%) and mesodermal (*e.g.* lateral plate mesoderm: 0.2 → 10.4%) cell types (**Fig. 1b**). This transition phenotypically coincides with visible cyst formation in the EBs. In contrast, phenotypic and transcriptional changes between day 14 and 21 appear less pronounced. We selected day 21 as the timepoint to harvest EBs in the context of our perturbation studies, due to its apparent stability and the most balanced distribution of different cell fates; we deemed the latter as particularly crucial to our statistical power for quantifying effects of a complex perturbation library in as many cell types as possible.

**Figure 1.**
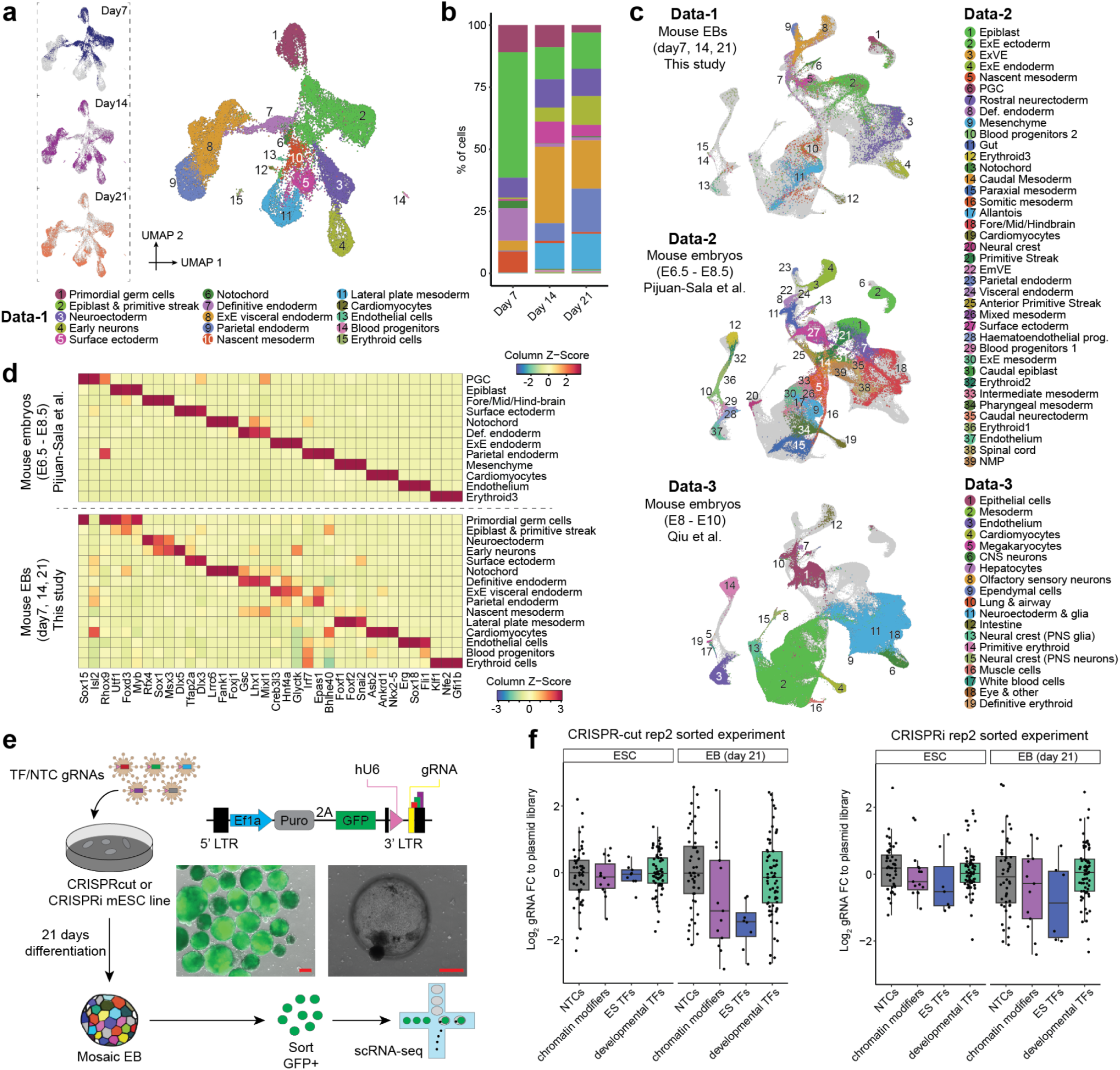
Characterizing embryoid bodies (EBs) as a model system for studying TF function in early development. **a,** 2D UMAP visualization of transcriptomes of 25,937 cells sampled from ∼150 pooled mouse EBs, with profiles captured at day 7, 14, and 21. Colors and numbers correspond to 15 cell cluster annotations as listed on the right, based on known marker genes (**Supplementary Table 1**). The same UMAP is shown three times on the left of the panel, with colors highlighting cells from different days. ExE: extraembryonic. **b,** Cell cluster composition of mouse EBs from each day. **c,** UMAP visualization of co-embedded cells from mouse EBs and mouse embryos^23,24^ at various developmental stages after batch correction of scRNA-seq data. The same UMAP is shown three times, with colors highlighting cells from either mouse EBs (top), embryos during gastrulation (middle), or embryos during early somitogenesis (bottom). **d,** Twelve cell types from mouse embryos were manually selected based on top marker genes and cell-type correlation analysis, as shown in **Supplementary Fig. 1a**, that matched cell types identified in mouse EBs. The top 3 TF markers of the 12 cell types were identified using the FindAllMarkers function of Seurat/v3^26^. The expression profiles of these TF markers were examined across various cell types within embryos during gastrulation^23^ (top) or mouse EBs (bottom). Each heatmap illustrates the mean gene expression values within each cluster, calculated from original UMI counts normalized to total UMIs per cell, followed by natural-log transformation. Cell type-specific TF expression is shared between gastrulating mouse embryos and mouse EBs. **e,** Experimental overview. To explore the function of individual TFs in cell fate determination, a monoclonal CRISPRcut or CRISPRi mouse ES line was transduced at low multiplicity of infection with a lentiviral CROP-seq library of gRNAs targeting TFs (3 gRNAs each) and non-targeting controls (NTCs) and containing GFP and puromycin selection markers. After optional puromycin selection to titrate the fraction of perturbed cells, ES cells were self-aggregated into mosaic EBs. After 21 days, GFP-positive cells were subjected to scRNA-seq and gRNA enrichment PCR. Representative microscopy images depicting a pool of GFP-positive EBs made from a puro-selected ES cell population (left) as well as a day 20 EB (right) are shown. The scale bar represents 200 µm. **f,** Changes in gRNA frequency for different target gene classes. The log2-fold-change in the proportion of gRNA guides was compared between mESCs (amplicon sequencing of gRNA from genomic DNA) and plasmid library, or between EBs (scRNA-seq on 10X Genomics platform) and plasmid library, in both the CRISPRcut pilot experiment (left) and the CRISPRi pilot experiment (right). The targets of the 144 gRNAs were classified into four groups: non-targeting controls (NTC), chromatin modifiers, ES TFs, and developmental TFs. Boxplots represent IQR (25th, 50th, 75th percentile) with whiskers representing 1.5× IQR.

**Table 1.** Overview of EB perturbation screens.

| Perturbation experiment name | CRISPR version | gRNA construct | gRNA targets | EB protocol | # of EBs profiled | Single cell protocol | gRNA detection | Sequencing | # of cells after QC (total / with gRNA) |
| --- | --- | --- | --- | --- | --- | --- | --- | --- | --- |
| wildtype time-series | none | none | none | mosaic | ~150 | 10X Gene Expression 3' v2 kit | none | NextSeq 500, R1: 26 bp, I1: 8 bp, I2: 0 bp, R2: 57 bp | 25,937 / 0 |
| pilot-mosaic | CRISPR cut and CRISPRi | CROP-seq GFP | 31 TFs (3 gRNAs each) + 51 NTCs (Table S3) | mosaic | ~1,500 | 10X Gene Expression 3' v3 | amplification from 10X cDNA library | NextSeq500, R1: 28 bp, I1: 8 bp, I2: 0 bp, R2: 55 bp | 58,037 / 32,188 |
| allTF-mosaic | CRISPR cut | CROP-seq GFP | 1,644 TFs (3 gRNAs each) + 999 NTCs (Table S3) | mosaic | ~2,200 | 10X Gene Expression 3' v3 | amplification from 10X cDNA library | NovaSeq 6000, R1: 28 bp, R2: 80 bp, I1: 10 bp | 190,387 / 140,129 |
| 125TF-mosaic | CRISPR cut | CROP-seq GFP | 125 TFs (3 gRNAs each) + 81 NTCs (Table S3) | mosaic | ~700 | 10X Gene Expression 3' v3 | amplification from 10X cDNA library | NextSeq 2000, R1: 28 bp, R2: 91 bp, I1: 8 bp and NovaSeq 6000, R1: 28 bp, R2: 91 bp, I1: 10 bp | 235,780 / 161,541 |
| pilot-monoclonal | CRISPRi | PiggyFlex v1 | 5,339 NTCs (Table S3) | monoclonal | 15 | 10X Gene Expression 3' v3 | 10X direct capture + organoid BC (v1, 15N randomer) amplification from 10X cDNA library | NextSeq 2000, Transcriptome : R1: 28 bp, I1: 8 bp, I2: 0 bp, R2: 91 bp. Organoid BC: R1: 28 bp, I1: 10 bp, I2: 0 bp, R2: 53 bp | 19,059 / 15,870 |
| 9TF-monoclonal -arrayed | CRISPRi | PiggyFlex v1 | 9 TFs (3 gRNAs each) + 3 NTCs (Table S3) | monoclonal | 100 | 10X Gene Expression 3' HT v3.1 | 10X direct capture + organoid BC (v2, 8N fixed-9N randomer) amplification from 10X cDNA library | NextSeq 2000, R1: 28 bp, I1: 10 bp, I2: 10 bp, R2: 90 bp | 69,598 / 55,063 |
| 8TF-monoclonal -pooled | CRISPRi | PiggyFlex v2 | 8 TFs (3 gRNAs each) + 3 NTCs (Table S3) | monoclonal | ~150 | 10X Gene Expression 3' HT v3.1 | 10X direct capture + organoid BC (v2, 8N fixed-9N randomer) amplification from 10X cDNA library | NextSeq 2000, R1: 28 bp, I1: 10 bp, I2: 10 bp, R2: 90 bp | 32,522 / 19,211 |
| 9TF-monoclonal -sci (same cell input as '9TF-monoclonal -arrayed' and '8TF-monoclonal -pooled') | CRISPRi | PiggyFlex v2 | 9 TFs (3 gRNAs each) + 3 NTCs (Table S3) | monoclonal | ~250 | sci-RNA-seq-3 | organoid BC (v2, 8N fixed-9N randomer) | NextSeq 2000, R1: 34 bp, I1: 10 bp, I2: 10 bp, R2: 84 bp | 46,754 / 7,930 |

To assess how well EBs recapitulate the cell type diversity observed in early mouse development, we next compared their single cell gene expression profiles to those from E6.5-8.5^23^ or E8-10^24^ mice. Co-embedding these datasets revealed that the decreasing epiblast population more closely resembles the epiblast and primitive streak population of younger mouse embryos, whereas the more differentiated cell types such as mesoderm, neuroectoderm, endoderm, cardiomyocytes, endothelial or blood cells share transcriptional similarities with their counterparts in both stages (**Fig. 1c**; **Supplementary Fig. 1a-b**). With the exception of neural crest, all major lineages in E6.5-E8.5 mice appear to have a corresponding cell type in EBs. In addition, some cell types specific to older mouse embryos, such as early neurons, were detected. Interestingly, the similarity of EBs to gastruloids^25^ is much more modest by comparison, mainly limited to PGCs, epiblast and primitive streak (**Supplementary Fig. 1c**).

While precisely quantifying similarities of cell types across datasets in high-dimensional space is challenging, here we were primarily interested in ascertaining to what extent cell type-specific expression patterns of TFs are recapitulated in mouse EBs, since this is the functionality that we sought to experimentally interrogate. Indeed, when we select the top three cell type-specific TFs for each cell type in the developing mouse, we found that those TFs are often specifically expressed in the cognate cell type in EBs, and vice versa (**Fig. 1d**; **Supplementary Fig. 1d**). Taken together, our results suggest that mouse EBs contain analogues of most major cell fates in early mouse development, and these exhibit a similar expression pattern of key transcriptional regulators.

### Setting up for large-scale CRISPR screens in mouse EBs

We next generated monoclonal mESC lines stably expressing CRISPR/Cas9 or CRISPRi^27^. To validate the capacity of these cells to form EBs and deplete target gene expression, we first transduced them with a pilot lentiviral CROP-seq^28^ library for which we had strong priors with respect to their expected effects (hereafter referred to as the ‘**pilot-mosaic**’ experiment; **Table 1**). Specifically, we targeted 31 TFs with three gRNAs each, including TFs thought to be essential in mESCs, chromatin modifiers thought to be dispensable in mESCs but important for EB differentiation, TFs previously associated with development of the three germ layers, and non-targeting controls (NTCs) (**Supplementary Fig. 2a**; **Supplementary Table 2-3**). By inserting a GFP into the CROP-seq gRNA vector backbone, we can quantify the transduction rate in mESCs and monitor for silencing of the gRNA during EB differentiation, and also select gRNA-expressing cells for downstream assays (**Fig. 1e**). Despite being controlled by the EF1α promoter, previously reported to be resistant to silencing during mESC differentiation^29,30^, only ∼20% of originally positive cells retained GFP expression at day 21 (**Supplementary Fig. 2b**).

We next sought to generate EBs with a low proportion of perturbed cells, simply by transducing them with gRNA-bearing lentivirus at a low MOI, and performing no selection. Performing scRNA-seq on the resulting EBs, we found that unsorted EB cells from CRISPRi or CRISPR/Cas9 clones (∼99% unperturbed, as determined by GFP FACS analysis) resemble cells from parental wildtype EBs in terms of their transcriptomes and cell type distribution (**Supplementary Fig. 2c-e**). In contrast, GFP+ cells from these same EBs (∼100% perturbed) were strongly enriched for less differentiated cell types (**Supplementary Fig. 2c-e**), consistent with expectation given the essential nature of many of the target genes in mESCs or EBs. Of note, this post-sorting enrichment could be caused by either the effects of perturbations on ESCs’ differentiation potential, or alternatively by an increase in lentiviral silencing of the CROP-seq construct in more differentiated cell types.

Increasing the proportion of perturbed cells in EBs from 1% (replicate 1) to 10-20% (replicate 2) by increasing transduction rates did not overtly impact EB growth, but did substantially increase the recovered number of gRNA-expressing cells (**Supplementary Fig. 2c-f**). After quality control, the average number of cells per target gene was 482 for CRISPRcut and 668 for CRISPRi. Comparing transcriptomes of cells with targeting guides to NTCs revealed on-target knockdown for most targets with CRISPRi. This was less consistently the case for CRISPRcut, which generates indels both with and without frameshifts, in line with previous comparisons of these modalities^31^ (**Supplementary Fig. 2g**).

Changes in overall gRNA representation compared to the plasmid library would imply that targeting the respective TFs affects cell survival and/or proliferation. As expected, we observed depletion of gRNAs targeting ESC-essential TFs in both stem cells (CRISPRi) and EBs (CRISPRi, CRISPRcut), whereas gRNAs targeting chromatin modifiers only became depleted upon EB formation (CRISPRi, CRISPRcut) (**Fig. 1f; Supplementary Fig. 2h-j**). We generally did not observe overall significant enrichment or depletion of the gRNAs targeting developmental TFs, possibly because their effects are limited to specific cell types in EBs. Apart from these qualitative assessments, the data was quite noisy, in part due to low gRNA assignment rates and differing transduction rates, two issues we resolved in future experiments.

Taken together, these pilot experiments: 1) confirm that genomic integration and expression of the CRISPR machinery does not overtly impact EB differentiation; 2) demonstrate our ability to measure perturbation identities and transcriptomes with high sensitivity in spite of lentiviral silencing phenomena; and 3) show that gRNAs targeting genes essential for EB formation show the expected drop-out effect in a pooled experiment. Of note, however, we also observed substantial variation in the effects of NTC gRNA controls (**Fig. 1f**), which in retrospect foreshadowed issues with reproducibility that we would encounter in our subsequent experiments.

### Performing large-scale TF perturbation screens in mosaic EBs

After establishing functional CRISPR machinery in our experimental system, we sought to perturb <u>all</u> mouse TFs and measure the impact on cell type composition in mosaic EBs (hereafter referred to as the ‘**allTF-mosaic**’ experiment; **Table 1**). We first generated a library consisting of gRNAs targeting all 1644 DNA or RNA-binding genes in the genome^32^, as well as 20% NTCs (3 gRNAs per target, for 5,931 gRNAs total; **Supplementary Table 3**). Since we expect most of these genes to be dispensable for EB formation, we transduced CRISPRcut and CRISPRi expressing stem cells such that cells receive at most one gRNA and selected successfully transduced cells with puromycin, resulting in a cell population of exclusively gRNA-expressing cells (**Supplementary Fig. 3a**). We observed stronger depletion of gRNAs targeting essential genes in the CRISPRcut than the CRISPRi line (**Supplementary Table 4**), and thus proceeded solely with the CRISPRcut perturbation modality. No obvious growth effects were observed upon pooled EB differentiation.

After dissociation of ∼1200 EBs at day 21, GFP-positive cells that had not undergone lentiviral silencing were sorted. We profiled the transcriptomes of 190,387 cells on the 10X Genomics platform, and recovered the associated gRNA for 140,129 (74%) of these (**Fig. 2a**; **Supplementary Fig. 3b**). On average, we recovered 75 cells per perturbation target across 1,612 targets (**Supplementary Fig. 3c**). As observed in the pilot, cell type composition revealed an increase in less differentiated cell types compared to WT EBs. However, stable NTC percentages across cell types imply this is due to increased silencing in more differentiated cells, rather than to perturbation effects (**Supplementary Fig. 3d-e**).

**Figure 2.**
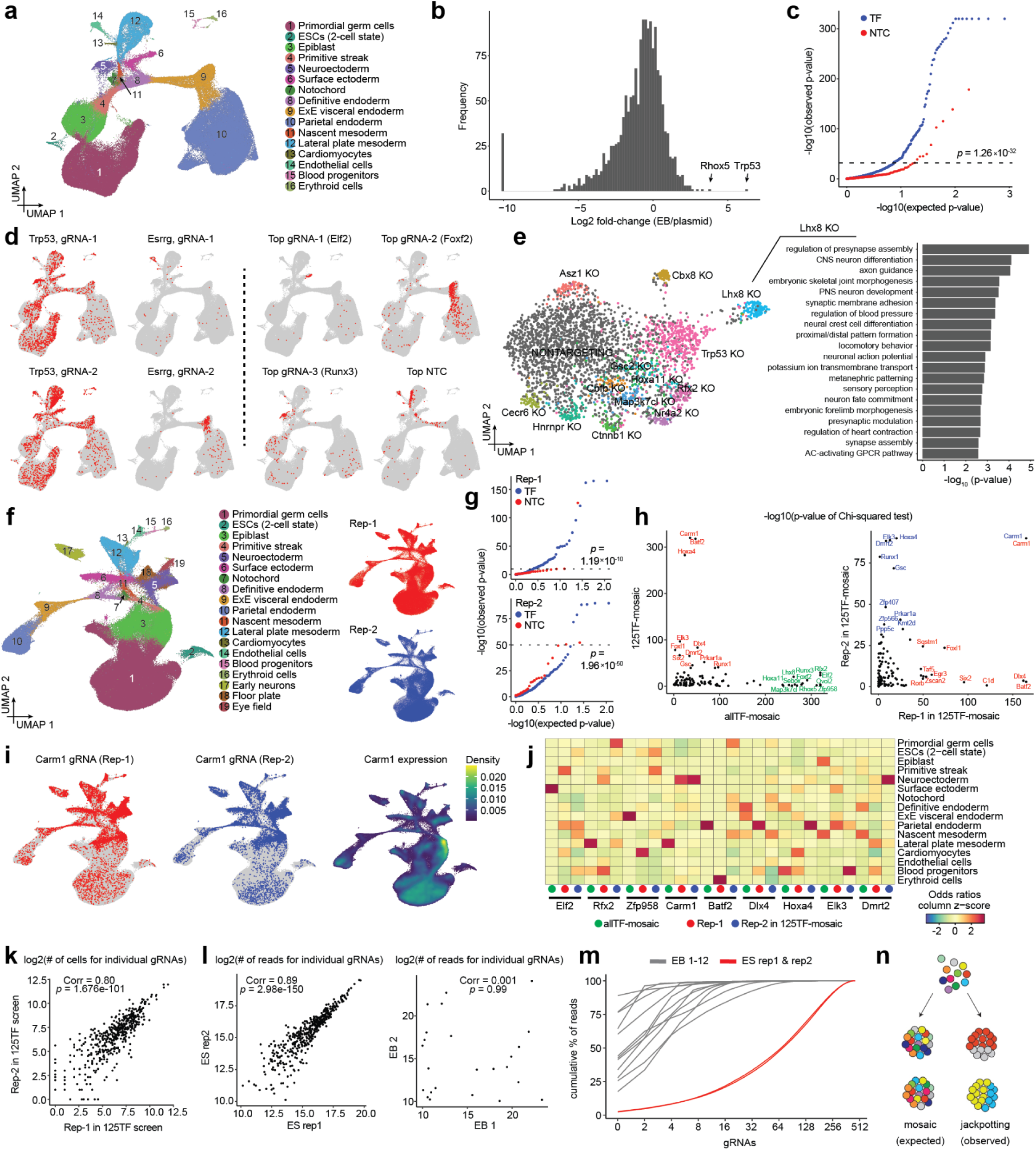
Performing large-scale TF screens in mosaic EBs. **a,** 2D UMAP visualization of 190,387 cells derived from ∼1200 pooled mosaic mouse EBs with CRISPR perturbation targeting 1,644 TFs (’allTF-mosaic’ experiment). Colors and numbers correspond to 16 cell cluster annotations as listed on the right. **b,** Changes in gRNA frequency for different target TFs. The log2-fold-change in the proportion of gRNA guides was compared between mouse EBs and plasmid library (’allTF-mosaic’ experiment). Target genes for gRNAs that drop out completely (left; visualized with small pseudocount) are listed in **Supplementary Table 5**. **c,** Quantile-quantile plot summarizing results of the ’allTF-mosaic’ experiment. TF targets with gRNAs detected in fewer than 50 cells or NTC gRNAs detected in fewer than 20 cells were filtered out. The remaining NTC gRNAs were randomly split into groups, each containing three gRNAs, after which cells within each group were randomly downsampled to 100 cells to approximately match the number of cells for each TF target. Chi-squared tests were conducted to compare cell type compositions between cells detected with each TF target (blue) or NTC gRNA group (red), and all cells detected with NTC gRNAs. The horizontal line represents the cutoff at the 0.05 quantile of the p-values calculated from the NTC gRNA group, and TF targets with p-values below this cutoff were considered significant. **d,** The same UMAP as in panel **a** is shown multiple times, with colors highlighting cells with the selected gRNAs detected. Left: Example of two gRNAs for the same target (*Trp53*) showing reproducible distribution, and two gRNAs for the same target (*Esrrg)* showing irreproducible distribution. Right: Top three significant TF gRNAs and the top NTC gRNA, ranked by p-values from the chi-squared tests. **e,** 2D UMAP visualization of 3,335 cells derived from lateral plate mesoderm cells (’allTF-mosaic’ experiment). The cells are colored and labeled according to their specific TF targets or NTC assignments. Only cells assigned to a single TF target are included in this analysis, and TF targets with fewer than 50 cells are excluded. Of note, the UMAP was generated using LDA as a dimensionality reduction method to visualize perturbation-specific clusters, utilizing the Mixscape library implemented in Seurat/v5^37^ to remove cells that have likely escaped perturbation^38^. Differentially expressed genes (DEGs) were identified between cells in which *Lhx8* was targeted vs. NTCs, and the top 20 terms upon GO ontology analysis are presented. **f,** 2D UMAP visualization of 235,780 cells derived from pooled mosaic mouse EBs with CRISPR perturbation targeting 125 selected TFs (’125lTF-mosaic’ experiment). Colors and numbers correspond to 19 cell cluster annotations as listed on the right. The same UMAP is shown twice on the right, with colors highlighting cells from each of the two biological replicates within the dataset. **g,** Quantile-quantile plots summarizing results of each replicate of ’125TF-mosaic’ experiment, with similar filtering as described for panel **c**. Chi-squared tests were conducted to compare cell type compositions between cells detected with each TF target (blue) or NTC gRNA group (red), and all cells detected with NTC gRNAs. The horizontal line represents the cutoff at the 0.05 quantile of the p-values calculated from the NTC gRNA group, and TF targets with p-values below this cutoff were considered significant. **h,** Comparison of results of chi-squared tests for each TF target for different experiments. Left: comparison of ‘allTF-mosaic’ vs. combined replicates of ‘125TF-mosaic’ experiments. Right: comparison of two replicates of ‘125TF-mosaic’ experiment. Selected TF targets with the top 10% of significant p-values from each dataset are labeled. **i,** The same UMAP as in panel **f** is shown multiple times, with colors highlighting cells with gRNAs targeting *Carm1* detected in replicate 1 (left) or replicate 2 (middle), or *Carm1* gene expression (right). Gene expression was visualized as a gene-weighted 2D kernel density plot using the Nebulosa package. **j,** The three most significant TF targets were selected for each of the three experiments based on the chi-squared tests: *Elf2*, *Rfx2*, and *Zfp985* (‘allTF-mosaic’ experiment); *Carm1*, *Batf2*, and *Dlx4* (replicate 1 of ‘125TF-mosaic’ experiment); and *Hoxa4*, *Elk3*, and *Dmrt2* (replicate 1 of ‘125TF-mosaic’ experiment). Early neurons, floor plate, and eye field in the ’125TF-mosaic’ dataset were merged into neuroectoderm to align with cell types annotated in the ‘allTF-mosaic’ experiment. For each TF target and each of the 16 cell clusters, the odds ratio was computed based on the frequency of cells within or outside the cell cluster between cells with the TF target and all cells with NTC gRNAs. **k,** The number of cells identified with each gRNA was compared between the two replicates of the ‘125TF-mosaic’ experiment. **l,** An independent experiment was performed to compare the gRNA abundances between two mESC replicates (left) or two of the twelve EB replicates (right). **m,** For each mESC or EB replicate from the independent experiment, the cumulative percentage of reads was plotted across gRNAs, ranked by their abundance from highest to lowest. The x-axis is displayed on a log2 scale, with tick labels showing the original (natural) values. **n,** EB differentiation led to a significant decrease in clonal complexity. Instead of the expected ∼100-1000 founding clones being equally represented in the final ∼10,000-30,000 cells (‘mosaic’), we observe stochastic clonal skewing resulting in ∼ 2-10 dominant clones (‘jackpotting’) .

When comparing relative gRNA frequencies in the day 21 EBs to gRNA frequencies in the original plasmid library, we observed expected changes in relative abundance, confirming successful CRISPR perturbation (**Fig. 2b**; **Supplementary Table 5**). For example, whereas known tumor suppressor gene *Trp53* exhibited an 80-fold increase due to proliferative advantage of cells in which this gene is knocked out, a subset of gRNAs were absent in EBs (corresponding to 32/1644 target genes). The targeted genes are mostly involved in essential processes such as DNA replication, transcription or translation (*e.g.* RNA polymerases, general transcription initiation factors, MCM complex, cell cycle regulators etc.).

In addition to impacts on ESC survival or EB formation, changes in relative gRNA frequency between cell types in the EB stage may reveal whether the target genes augment or inhibit the development of certain cell types. We quantified differential distribution of gRNAs relative to NTCs across cell types with a chi-square test, by combining all three gRNAs targeting a certain gene and comparing their distribution to a similar number of cells containing sets of three individual NTC gRNAs, to mimic targeting guides as closely as possible. This analysis identified a substantial number of gRNAs as differentially distributed across cell types compared to NTCs (13.4% with an empirical FDR of 0.05). Some of these were expected, but there were also many unanticipated enrichments and depletions (**Fig. 2c-d**).

Since cell type definitions rely on empirically chosen cluster resolution and might mask more local enrichments or depletions within larger clusters, we also quantified differences in local KNN-graph neighborhoods in the high-dimensional PCA space (lochNESS score^33^). While this measure is agnostic to cell type resolution and robust across different chosen k-parameters, it preferentially identifies local enrichments rather than depletions due to the relatively low cell number per gRNA in this experiment. As expected, cell-type specific chi-square test and cell-type-agnostic lochNESS scores are correlated (**Supplementary Fig. 3f**), although both measures preferentially detect different patterns: Whereas lochNESS appears better at identifying local enrichments within larger cell types (*e.g. Foxp1* targeting is locally enriched in a specific region of the endoderm), the chi-square test appears better at identifying broad germ-layer level depletions (*e.g. Trp53* targeting is depleted in parietal endoderm) (**Supplementary Fig. 3f-g**).

In addition to relative gRNA abundance across cell types, for targets with sufficient cell numbers, we can also detect transcriptional changes that occur within cell types as a result of the perturbations. For example, within the mesoderm, we observed downregulation of apoptosis-related genes in *Trp53* KO cells. Also in mesoderm, we observed upregulation of osteogenic and neurogenic genes in *Lhx8* KO cells (**Fig. 2e; Supplementary Table 6**). On the other hand, the low number of EBs sampled, the low cell number per individual gRNA, and the preferential depletion of gRNAs targeting genes present in the pilot screen, made it challenging to comprehensively assess reproducibility of effects either across gRNAs in the same experiment or separate experiments. This is crucial to gain confidence in the results, especially in light of discordant enrichment patterns observed for some guides with the same targets and significant differential distribution observed for some NTCs (**Fig. 2c-d**).

### Confounding effects of gRNA genotype imbalances on mosaic perturbation screens

To assess reproducibility of the observed cell type-specific gRNA distributions, we selected 125 significantly differentially distributed targets from the ‘allTF-mosaic’ screen for a new CRISPRcut experiment (hereafter referred to as the ‘**125TF-mosaic**’ experiment; **Table 1**; **Supplementary Table 3**). We then collected a larger number of cells per target and performed two independent biological replicates by transducing this library into two separate stem cell populations and inducing independent EB differentiations. This resulted in an average of 205 and 261 cells per gRNA, or 557 and 708 cells per target, for the two replicates (**Fig. 2f**; **Supplementary Fig. 3b-c**). Strikingly, when comparing cell type distribution of gRNAs using the chi-square test, we found that although we again detect many significantly differentially distributed targets, these were <u>not</u> reproducible across experiments, with the notable exception of *Carm1* (**Fig. 2g-i**; **Supplementary Fig. 3h**). This target (*Carm1*) incidentally had the highest cell number within the screen and its targeting gRNAs were reproducibly depleted in the epiblast, where it was most highly expressed. In contrast, they were enriched in differentiated cell types, in line with Carm1’s key role in maintaining pluripotency by regulating Oct4/Sox2 expression^34^. Essentially all other targets exhibited highly significant but irreproducible enrichments across different cell types (**Fig. 2j**) and we again observed p-value inflation for NTCs, especially for the second replicate (**Fig. 2g**). One possible explanation for this irreproducibility could be bottlenecking of the overall gRNA library complexity during ES or EB culture, leading to differences in overall representation. However, this did not appear to be the case, as overall gRNA frequencies in EBs were highly correlated between the two biological replicates (**Fig. 2k**).

An alternative explanation is bottlenecking of gRNA genotype frequencies in individual mosaic EBs, *i.e.* if cells of a certain type and bearing a certain gRNA become overrepresented not due to the effect of their perturbation, but rather due to stochastic events during mosaic EB development. It has been suggested that EBs arise from aggregates of hundreds to thousands of ESCs^35^, which continue dividing for roughly a week before reaching a postmitotic state. To quantitatively test this assumption, we hand-picked individual day 21 EBs and counted their final cell number, which consisted of around 10,000-30,000 cells. After verifying our ability to accurately detect gRNA genotype frequency from genomic DNA across similar cell numbers in mESCs, we quantified gRNA genotype frequencies within 12 individual, hand-picked EBs (**Fig. 2l**; **Supplementary Fig. 3i**).

This revealed a surprising distribution of gRNA genotype frequencies: although these EBs were mosaic as expected, they contained fewer than 100 distinct gRNAs and were strongly dominated by 2-10 of these that were 2-3 orders of magnitude more prevalent than the rest (**Fig. 2m**). The identities of these ’jackpot’ gRNAs were highly divergent across individual EBs. Accordingly, based on the number of EBs sampled in this experiment (∼350 EBs per replicate, ∼500-700 cells per target with most EBs dominated by 2-4 gRNAs), we estimate that most cells for a given target are likely derived from only 2-4 EBs, which is not sufficient to average out inter-EB heterogeneity with respect to cell type composition^36^. Put another way, this stochastic, highly non-uniform contribution of clones within EBs dramatically reduces our power, explaining not only the differential cell type distributions seemingly associated with TF-targeting gRNAs in our experiment, but also their irreproducibility (**Fig. 2n**).

### A protocol for generating monoclonally-derived, individually-barcoded embryoid bodies

To summarize, performing perturbation scRNA-seq screens of mosaic EBs led us to the following conclusions: 1) Mosaic EBs have markedly lower clonal complexity than expected given the number of founding cells; 2) Detectable clones are highly skewed with respect to their frequencies, with fewer than 10 clones composing the vast majority of a given EB; 3) This jackpotting appears to be largely stochastic rather than driven by perturbation-specific effects; 4) While there is some underlying biological signal, as evidenced by higher significance of targeting guides than NTCs in most experiments and a subset of reproducible enrichments across guides and replicates that rise above the noise, these effects appear to be mainly detectable when cell numbers per target are high and one is averaging over many EBs (*e.g.* p53 or Carm1); and 5) In addition to stochastic jackpotting, a further factor limiting our power was the low number of cells recovered per perturbation, due to silencing of the gRNA and associated GFP during differentiation.

Towards resolving or at least mitigating these issues, we set out to develop a new system for EB perturbation with two crucial differences. First, to increase the number of cells recovered per perturbation, we sought to switch from the lentiviral backbone vector, which is prone to silencing^39^, to the piggyBac transposase-transposon system (**Fig. 3a**). Second, we sought to develop a protocol for generating <u>monoclonal</u> rather than mosaic EBs, *i.e.* wherein each EB originates from a single mESC. By definition, monoclonal EBs would not suffer from bottleneck-induced differences in gRNA frequency across cell types, since all cells within the same EB would be expressing the same gRNA (**Fig. 3b**). In addition, monoclonality would permit adding an ’organoid barcode’ to each mESC-of-origin and thus its descendants, enabling quantification of differences between cells that received the same perturbation and were differentiated in the same dish, yet were derived from different monoclonal EBs.

**Figure 3.**
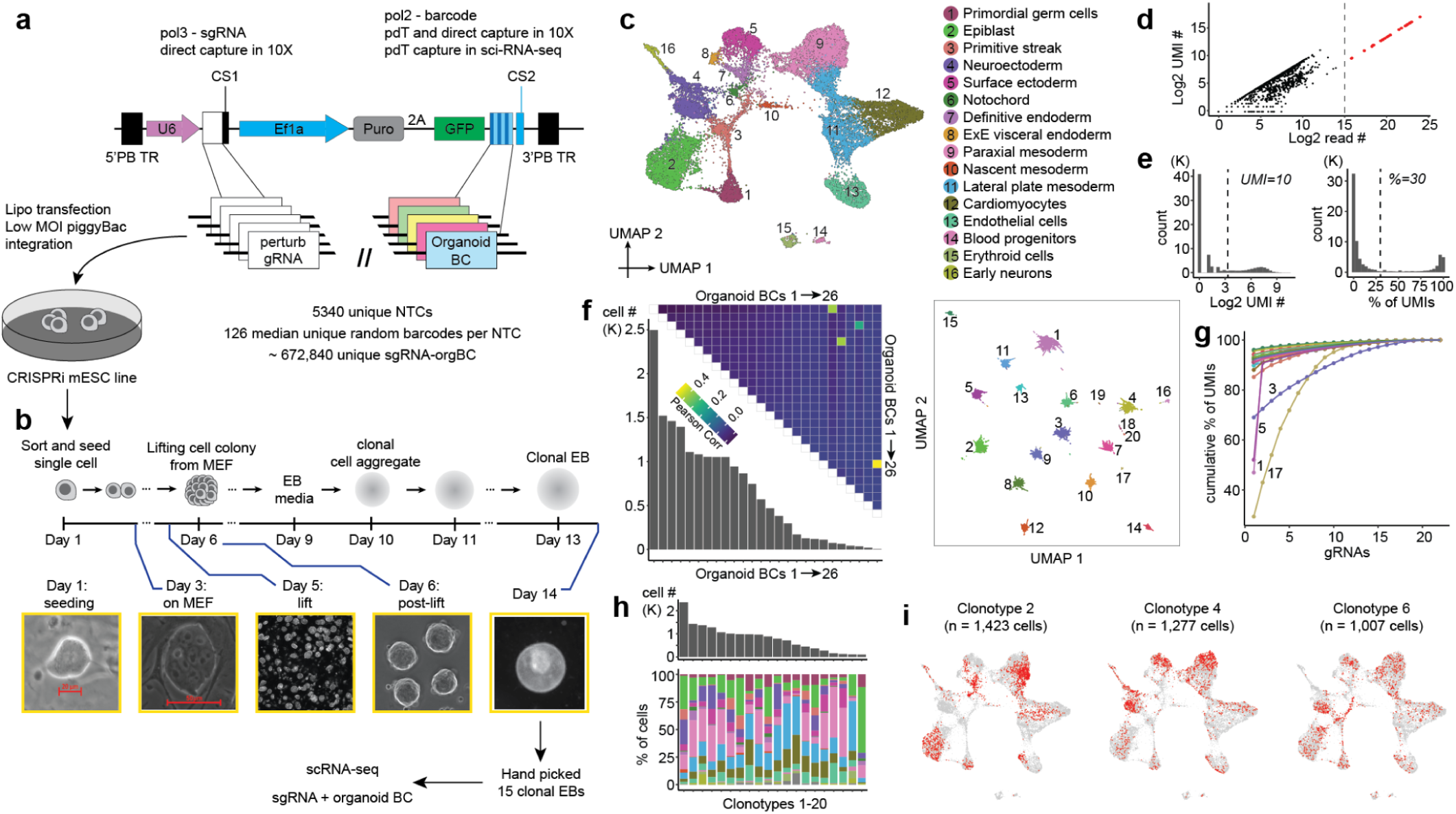
Development of a monoclonal EB screening platform. **a,** Schematic of piggyFlex construct and its application to co-express a gRNA as well as an organoid BC that is informative with respect to both gRNA identity and organoid identity. **b,** Schematic of proof-of-concept experiment for generating and validating monoclonal EBs, each founded by an mESC that is stably tagged with gRNA/organoid BC-expressing piggyFlex construct. In this experiment, all gRNAs were non-targeting controls. To obtain a pool of monoclonal EBs, piggyFlex mESCs were sorted and seeded at low density on mouse embryonic fibroblasts. Individual mESCs grew into colonies for 5 days followed by lifting with Collagenase IV treatment and gentle agitation. Colonies, now clonal aggregates, were then differentiated into EBs on low adherent plates for 8 days. **c,** 2D UMAP visualization of 19,059 cells derived from 15 manually selected, pooled monoclonal mouse EBs at day 8. Colors and numbers correspond to 16 cell cluster annotations. **d,** The log2-scaled UMI count and log2-scaled read count are plotted for individual organoid BCs. For the subsequent analyses, 26 organoid BCs with a log2-scaled read count >= 15 were selected. **e,** For each of these 26 organoid BCs, the log2-scaled UMI count within each cell (left) and the proportion of UMI counts within each cell (right) are plotted. Organoid BCs were assigned to individual cells if the UMI count >= 10 and the proportion >= 30%. **f,** After excluding cells without any assigned organoid BCs, the UMI count across the 26 organoid BCs for individual cells (*n* = 16,512) was normalized by the total count per cell, followed by scaling the normalized counts per organoid BC. Pairwise Pearson correlations were then computed for each pair of organoid BCs, are displayed in the top heatmap. The number of cells assigned to each organoid BC is shown in the bottom plot. On the right, dimension reduction using PCA was performed on the UMI counts of organoid BCs across 16,512 cells, followed by visualization in a 2D UMAP. After that, cells assigned multiple organoid BCs were excluded, except for those with correlation coefficients >0.1, which were considered for merging. Organoid BC pairs 1 & 21, 3 & 24, 5 & 22, and 20 & 26 were merged. Cells were then grouped based on their assigned unique organoid BC or merged organoid BCs, resulting in a total of 20 clonotypes comprising 15,870 cells, after excluding clonotypes with fewer than 50 cells (organoid BCs 23 & 25). In the UMAP, cells are colored and labeled by their assigned clonotypes, with unassigned cells colored in gray. **g,** After filtering out gRNAs expressed in <1% of cells, 22 gRNAs were retained. Cells with fewer than 10 UMIs of gRNAs were excluded, followed by normalizing the gRNA UMI counts to the total count per cell. The proportion of UMI counts per gRNA was calculated for individual cells, and the average proportions were computed for cells within each clonotype. Subsequently, the cumulative proportion of UMIs across gRNAs, ranked by their abundance from highest to lowest, is plotted for each of the 20 clonotypes. Clonotypes with multiple assigned gRNAs are labeled. **h,** Cell type composition of mouse EBs from each clonotype bin. Colors are defined in panel **c**. The top plot displays the number of cells sampled for each clonotype. **i,** The same UMAP as in panel **c** is shown multiple times, with colors highlighting cells from the three most abundant clonotypes, each dominated by a single assigned gRNA (>90% of total UMIs as shown in panel **g**).

To support these changes, we leveraged the piggyFlex vector^40^ (**Fig. 3a**). In this dual RNA expression piggyBac construct, a U6 (Pol3) promoter-driven gRNA is co-expressed with a EF1alpha (Pol2) promoter-driven GFP, a puromycin resistance gene and, as implemented here, a 3’ UTR organoid barcode (BC). The organoid BC is a sequence barcode that is informative with respect to which gRNA perturbation is encoded while also providing a unique identifier for a given monoclonal EB and distinguishing independent ’instances’ of the same gRNA perturbation. Barcode-gRNA associations were either pre-encoded during synthesis or determined by sequencing of the plasmid library, with optimized strategies to minimize rates of gRNA-barcode dissociation. The piggyFlex vector also includes capture sequences that enhance recovery of Pol3 gRNA transcripts during scRNA-seq^41^.

To validate piggyFlex, we integrated the construct into mESC genomes via co-transfection with the piggyBac transposase at a low MOI and monitored GFP expression at day 7, 14 and 21 of EB formation. On day 21, 88% of cells were GFP-positive (compared to 21% of cells sampled from EBs expressing GFP from the integrated lentiviral CROP-seq construct; **Supplementary Fig. 4a**), confirming that piggyFlex is more resistant to silencing in differentiating EBs, and eliminating the need for FACS prior to scRNA-seq. To verify that the CRISPR machinery is effective when expressed from this construct, we transfected mESCs with a library of NTCs and gRNAs targeting essential genes. As expected, gRNAs targeting essential genes were depleted after 14 days in culture, while gRNAs targeting NTCs were not (**Supplementary Fig. 4b**).

To generate monoclonal EBs, we transfected a complex piggyFlex library containing 5,338 NTC gRNAs, each of which was associated with randomly linked organoid BCs (mean 128 barcodes per NTC) into mESCs at a low MOI and selected the cells for puromycin resistance for 10 days (hereafter referred to as the ‘**pilot-monoclonal**’ experiment; **Table 1**; **Supplementary Table 3**). Sequencing of the plasmid library enabled us to assign each organoid BC to a specific gRNA, *i.e.* a lookup table. We then sorted single mESCs onto a mouse embryonic feeder (MEF) layer in 6-well plates at different densities, with the aim of growing monoclonal colonie (**Fig. 3b**). After 5-6 days, colonies were gently lifted with Collagenase IV treatment followed by agitation and transferred to non-adherent plates. The suspended clonal aggregates became spherical and continued to differentiate into EBs. A total of 15 monoclonal mouse EBs at day 8 were manually selected for scRNAseq on the 10X Genomics platform (**Fig. 3c**). Organoid BCs and gRNAs were subjected to enhanced recovery via enrichment PCR (organoid BCs) or direct capture feature barcoding (gRNAs). Integration of transcriptomes from monoclonal EBs with natural embryos and other *in vitro* models demonstrated robust germ layer capture, whereas integration with mosaic wildtype EBs revealed a relative increase in the mesodermal and a decrease in the endodermal populations (**Supplementary Fig. 5a-c**).

For cell-to-EB assignments, high-abundance organoid BCs (*n* = 26 with log2 total organoid BC read counts ≥15) were assigned to individual cells if the organoid BC UMI count was ≥10 and the proportion of organoid BC UMI counts was ≥30% within that cell (**Fig. 3d-e**). After excluding cells without assigned organoid BCs (∼13%), we calculated pairwise correlations of the organoid BCs. Cells assigned multiple organoid BCs (∼4%) were excluded, except those with organoid BC pairs showing correlation coefficients > 0.1, which were considered for merging (**Fig. 3f**). These likely correspond to EBs derived from cells which have multiple gRNAs integrated into their genomes. Cells were grouped based on their assigned unique or merged organoid BCs, resulting in 20 ‘clonotypes’ after excluding those with fewer than 50 cells. Of note, we expected to capture 15 clonotypes based on the 15 EBs that went into this experiment. The additional 5 clonotypes could be caused by either floating EB fragments contaminating the hand-picked EBs, some EBs starting from 2 cells instead of 1, or insufficient co-capture of organoid BCs in EBs derived from cells with multiple integrations.

Next we sought to assess whether organoid BCs captured from Pol2 transcripts were correctly revealing the identity of the linked perturbation encoded by a Pol3-driven gRNA. We can test this directly in this experiment, as gRNA sequences are co-captured via feature barcoding on the 10X Genomics platform. After filtering out gRNAs expressed in fewer than 1% of cells, 22 gRNAs were retained. Most clonotypes were dominated by a single gRNA, with 16 clonotypes exhibiting a gRNA UMI proportion >80% (**Fig. 3g**). Of the 31 organoid BC-gRNA combinations detected in the 20 clonotypes with gRNA UMI proportions >10%, 17 were identified using the lookup table derived from the initial sequencing of the plasmid library that associated organoid BCs with specific gRNAs. Of note, for later experiments described in this paper that leverage piggyFlex (**Fig. 4**) we switched to a predefined constant barcode subsequence as part of the organoid BC that is unequivocally linked to a specific gRNA (while retaining a degenerate subsequence as well to mark the individual EB), so as to circumvent the limitations of sequencing-based association.

**Figure 4.**
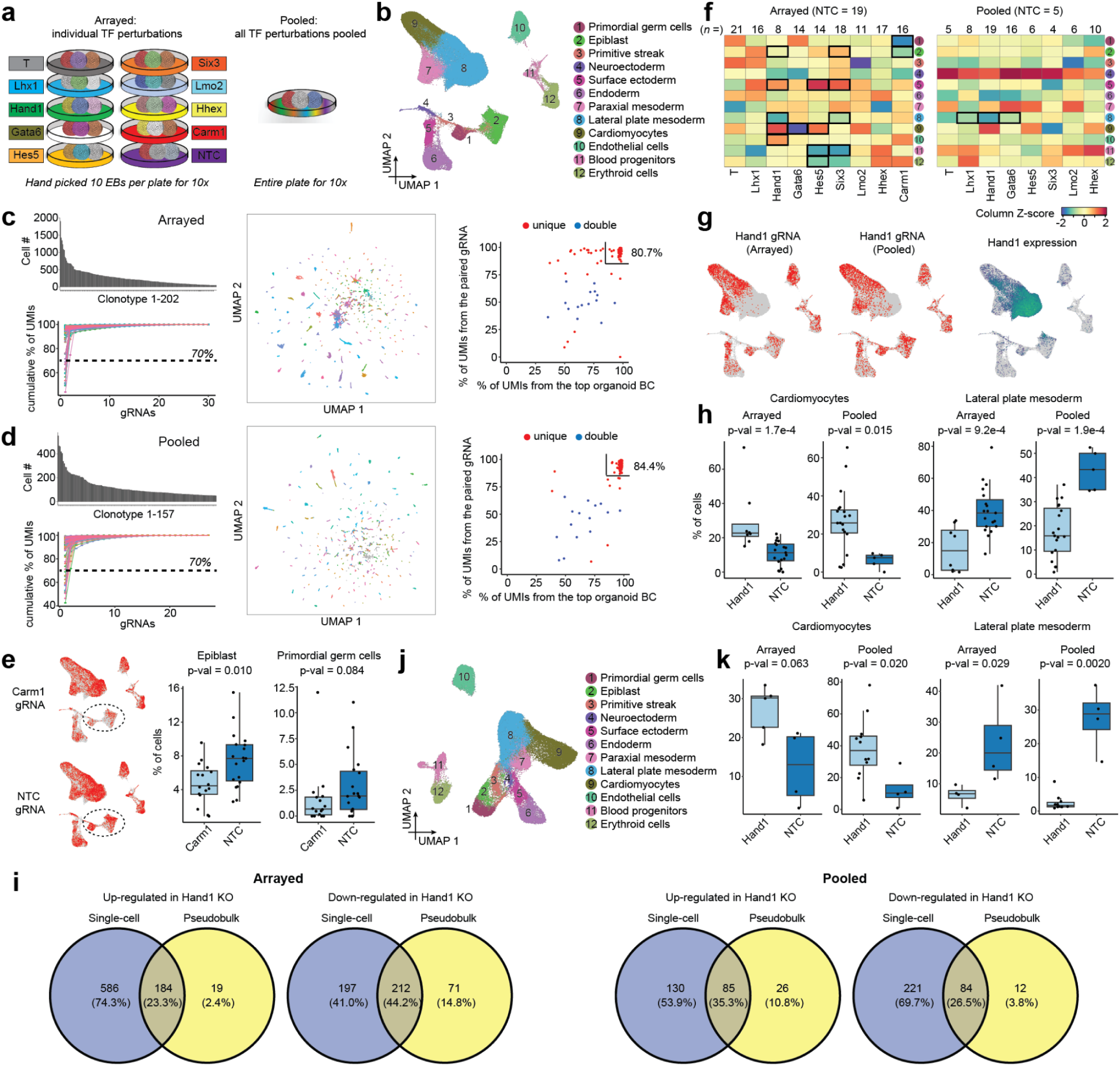
Proof-of-principle TF screen in monoclonal EBs. **a,** In the arrayed experiment (left), also referred to as the ‘9TF-monoclonal-arrayed’ experiment, nine TF perturbations and a non-targeting control (NTC) were evaluated using 10 handpicked monoclonal EBs per separate condition, each with unique gRNAs and organoid BCs, pooled and processed to scRNA-seq across 12 lanes of the 10X Genomics platform. In the pooled experiment (right), also referred to as the ‘8TF-monoclonal-pooled’, eight TF perturbations (Carm1 was excluded) and a non-targeting control (NTC) were assessed in a pooled screen of approximately 150 monoclonal EBs, pooled and processed to scRNA-seq across 4 lanes of the 10x Genomics platform. **b,** 2D UMAP visualization of 102,120 cells derived from pooled monoclonal mouse EBs from the two experiments. Colors and numbers correspond to 12 cell cluster annotations as listed on the right. **c,** In the arrayed experiment, a total of 202 clonotypes with at least 50 cells were identified. Top left: The cell count for each clonotype is shown. Bottom left: The cumulative proportion of UMIs across gRNAs, ranked by abundance from highest to lowest, is plotted for each of the 202 clonotypes. Middle: Dimensionality reduction using PCA was performed on the UMI counts of organoid BCs across 58,232 cells, followed by visualization in a 2D UMAP. Cells are colored and labeled by their assigned clonotypes, with unassigned cells colored in gray. Right: Proportions of UMIs from the most abundant organoid BC within each clonotype and the UMIs of the corresponding paired gRNA, as determined by the constant organoid BC part, in the directly captured gRNA library. Uniquely assigned clonotypes are shown in red, while doubly assigned ones are in blue. 80.7% of clonotypes exceed 85% on both axes, indicating strong agreement between gRNA/perturbation identities inferred from the organoid BCs vs. direct capture. Uniquely assigned clonotypes are shown in red, while doubly assigned ones are in blue. **d,** Same as described for panel **c**, for 157 clonotypes from the pooled experiment. In the middle panel, 22,897 cells are represented. In the right panel, 84.4% of clonotypes exceed 85% on both axes, again indicating strong agreement between gRNA/perturbation identities inferred from the organoid BCs vs. direct capture. **e,** The same UMAP as in panel **b** is shown, with colors highlighting cells from clonotypes with gRNAs targeting *Carm1* detected in the arrayed experiment. This can be quantified on an ‘per-individual-EB’ level using clonotypes (right, each dot corresponds to an individual clonotype). Cell fractions were compared between clonotypes assigned with *Carm1* gRNA vs. NTC gRNA from the arrayed experiment (*n* = 16 for Carm1 and 19 for NTC) for epiblast (left) and primordial germ cells (right), respectively. Wilcoxon tests were performed, and the resulting p-values are reported. Boxplots represent IQR (25th, 50th, 75th percentile) with whiskers representing 1.5× IQR. **f,** Hooke^42^ was used to compare cell abundances across 12 identified cell types between clonotypes assigned with each TF target vs. NTCs in the arrayed experiment (left) or pooled experiment (right). Clonotypes with fewer than 100 cells were excluded. For each cell type, the resulting natural-log-fold changes in cell abundances between clonotypes assigned to each TF target or NTCs are shown in the heatmap. Each column is scaled, and significant changes (FDR < 0.1) are highlighted by black rectangles. **g,** The same UMAP as in panel **b** is shown multiple times, with colors highlighting cells from clonotypes with gRNAs targeting Hand1 detected in the arrayed experiment (left), the pooled experiment (middle), and Hand1 gene expression (right). **h,** Cell fractions were compared between clonotypes bearing *Hand1* gRNAs vs. NTC gRNAs from the arrayed (*n* = 8 gRNAs for *Hand1* and 19 for NTC) and pooled experiments (*n* = 19 gRNAs for *Hand1* and 5 for NTC) for cardiomyocytes and lateral plate mesoderm, respectively. Wilcoxon tests were performed, and the resulting p-values are reported. Boxplots represent IQR (25th, 50th, 75th percentile) with whiskers representing 1.5× IQR. **i,** The number of significantly up- and down-regulated genes (adjusted p-value < 0.05) differentially expressed between Hand1 knockouts vs. NTC monoclonal EBs were determined using pseudobulk or single cell profiles from two experiments. For pseudobulk analysis, individual clonotypes assigned to *Hand1* gRNAs or NTC gRNAs were compared using DESeq2^47^ to identify differentially expressed genes. For single-cell analysis, cells were classified as bearing *Hand1* or NTC gRNAs based on UMI abundance exceeding 90%, and differentially expressed genes were identified using the *FindMarkers* function in Seurat^37^. **j,** 2D UMAP visualization of 46,754 cells from pooled monoclonal mouse EBs across two experiments, profiled with sci-RNA-seq3. Colors and numbers correspond to 12 cell cluster annotations as listed on the right. **k,** From the sci-RNA-seq3 data, cell fractions were compared between clonotypes bearing *Hand1* gRNAs vs. NTC gRNAs from the arrayed (*n* = 5 gRNAs for *Hand1* and 4 for NTC) and pooled experiments (*n* = 12 gRNAs for *Hand1* and 4 for NTC) for cardiomyocytes and lateral plate mesoderm, respectively. Wilcoxon tests were performed, and the resulting p-values are reported. Boxplots represent IQR (25th, 50th, 75th percentile) with whiskers representing 1.5× IQR.

Altogether, this pilot experiment establishes our ability to to generate monoclonal EBs, to circumvent gRNA silencing by switching to a piggyBac-based vector, and to concurrently capture both the perturbation and the ’individual’ EB identity with organoid BCs. Furthermore, assigning distinct clonotypes to cells facilitated comparison of cell type composition across individual EBs. Even though this experiment did not contain any active perturbations, substantial heterogeneity in cell-type compositions were detected across different clonotypes, reinforcing the ’individual-to-individual’, intra-batch heterogeneity of even unperturbed embryoid bodies (**Fig. 3h-i**). At these levels of heterogeneity, much higher cell coverage per EB or many individual EBs would be needed to reliably detect differences in cell type composition, relative to a situation in which EBs exhibited uniform composition.

### Proof-of-concept for a monoclonal EB pooled screening platform

As a proof-of-concept of a genetic screen of pooled, sequence-tagged monoclonal EBs, we focused on measuring the effects of piggyFlex-CRISPRi mediated perturbations of nine developmentally relevant TFs (*Carm1, Gata6, Hand1, Hes5, Hhex, Lhx1, Lmo2, Six3*, and *T*) on cell type composition.

As a control experiment (’arrayed’ in **Fig. 4a**; also referred to as the ‘**9TF-monoclonal-arrayed**’ experiment; **Table 1**), we transfected a series of dishes of mESCs with a small pool of piggyFlex constructs encoding three gRNAs each targeting the same TF (or three NTC gRNAs). Individual mESCs from each dish were then used to generate separate pools of monoclonal EBs, *i.e.* one dish per perturbation (or NTCs). Ten monoclonal EBs per perturbation (or NTCs) were then handpicked, pooled, dissociated to single cell suspensions and subjected to scRNA-seq (**Supplementary Table 3**). Importantly, even if founder mESCs and the resulting EBs bear multiple piggyFlex integrants, the arrayed workflow guarantees that these perturb the same target gene.

For our proof-of-concept experiment (’pooled’ in **Fig. 4a**; also referred to as the ‘**8TF-monoclonal-pooled**’ experiment; **Table 1**), we transfected a single dish of mESCs with a pool of piggyFlex constructs encoding 27 gRNAs (3 gRNAs x 8 TF, plus 3 NTC gRNAs) at a low MOI. After a selection step, these mESCs were used to generate monoclonal EBs, each tagged with an organoid BC that is informative with respect to both individual EB identity and the encoded perturbation (**Supplementary Table 3**). Of note, this experiment targets the same TFs as in the arrayed experiment while intentionally excluding targeting of *Carm1*, as we were concerned that mESCs bearing its perturbation might outcompete other mESCs in a pooled setting given our earlier observations. Instead of handpicking, approximately 150 monoclonal EBs were collected by aspiration, dissociated to single cell suspensions and subjected to scRNA-seq. A key point is that although the number of perturbations is quite limited for this proof-of-concept experiment, this pooled workflow is considerably more scalable to additional targets than the arrayed workflow, and primarily limited by scRNA-seq costs.

We next sought to compare the perturbation effects observed via the pooled vs. arrayed screens. Altogether, we profiled 32,522 scRNA-seq profiles from pooled EBs and 69,598 from arrayed EBs. We identified 12 cell types among these transcriptomes, whose annotations were consistent with our earlier experiments (**Fig. 4b**). Cell-to-EB assignments were performed separately for each experiment as previously. From the arrayed experiment, we identified 202 clonotypes across the 10 conditions comprising 55,063 cells (mean: 273 cells, median: 209 cells, range: 50–1,907 per clonotype) (**Fig. 4c**). This is considerably more than the expected 100 clonotypes, which could reflect too high of a seeding density resulting in some EBs being derived from more than one cell (as cells with multiple integrants are expected to be collapsed during clonotype assignment). From the pooled experiment, we identified 157 clonotypes comprising 19,211 cells (mean: 122 cells, median: 93 cells, range: 50–552 per clonotype), roughly in line with our expectation of 150 (**Fig. 4d**).

To assign gRNAs to clonotypes, UMI proportions across gRNAs directly captured from Pol3 transcripts were calculated for each cell and averaged within clonotypes. Clonotypes with a single gRNA averaging ≥70% were classified as ’unique’ assignments, while the top two most abundant gRNAs were retained for the remaining clonotypes as ’double’ assignments. In the arrayed experiment, 93% (n=187) clonotypes received unique gRNA assignments, and 7% (n=15) double gRNA assignments (in each case targeting the same gene, as expected for the arrayed experiment). In the pooled experiment, 89% (n=139) clonotypes received unique gRNA assignments, and 10% (n=15) double gRNA assignments (in all but one case targeting different genes, as expected for the pooled experiment), and 2% (n=3) were discarded due to poor gRNA recovery. Particularly for unique assignments, we observed strong concordance between gRNA assignments based on the organoid BC and the gRNAs captured in scRNA-seq data via feature barcoding (right panels in **Fig. 4c-d**), validating our assignment of perturbation identities solely from the organoid BC.

After assigning each cell to an EB clonotype and each clonotype to a perturbation, we examined changes in cell type composition in response to target gene perturbation across both experiments. For example, in the arrayed workflow, *Carm1* gRNAs were depleted in less differentiated cell states (**Fig. 4e**), in line with our earlier mosaic screen (**Fig. 2i**). However, in sharp contrast with the mosaic screen, monoclonal EBs crucially enable us to statistically assess the consistency of cell type composition across many individual EBs (boxplots in **Fig. 4e**), as well as to circumvent the possibility that perturbations with a growth advantage outcompete other perturbations within an EB.

To investigate differential cell abundances between each perturbation and wild type across the entire dataset, we applied a new computational method ’*Hooke*’, which uses Poisson-Lognormal models to account for variability in count data while assessing how perturbations impact cell types’ relative abundances^42^. In the arrayed experiment, eight cell types exhibited composition changes (FDR < 0.1) in response to five TF knockouts, compared to wild type (**Fig. 4f**; **Supplementary Table 7**). One of these changes was replicated at this FDR cutoff in the pooled experiment. Specifically, we observe reproducible depletion of the lateral plate mesoderm in *Hand1* gRNA-expressing EBs, as well as enrichment of cardiomyocytes (although the latter was not statistically significant in the pooled experiment). Whereas the essential role of Hand1 in lateral mesoderm development is well-established^43^, it was recently shown that *Hand1* deletion drives multipotent progenitors towards a cardiomyocyte cell fate^44^ (**Fig. 4g-h**; **Supplementary Table 8**). The small number of reproducible cell type changes and reduced number of significant changes in the pooled experiment is likely due to two factors. First, power issues -- *i.e.* the differences in the total number of cells profiled for the same number of expected clonotypes (in average, 273 vs. 122 cells per clonotype), the number of NTC clonotypes that can be used for the statistical test (19 vs. 5) and the relatively low number of clonotypes profiled per perturbation (154 vs. 79 clonotypes used for *Hooke* analysis). Second, individual-to-individual heterogeneity -- *i.e.* even in the context of the same perturbation (or NTC), monoclonal EBs exhibit considerable variation in cell type composition.

In addition to assessing cell type composition changes across perturbation conditions, high-content single cell screens have the potential to reveal which genes are affected within each cell type. Importantly, our monoclonal platform allows us to assess the reproducibility of differentially expressed genes across individual EBs, by pseudobulking single cell profiles per clonotype. For example, whereas standard single differential gene expression analyses using *Seurat*^37^ without clonotype information would nominate 770 differentially expressed genes in this experiment, an analysis that controls for clonotype reveals that 76% of these are potentially false positives driven by EB-to-EB heterogeneity rather than perturbations (**Fig. 4i**).

The variability across individual EBs suggests that a large number of EBs for each perturbation condition will be required for drawing robust conclusions both in regards to cell type composition and intra-cell type gene expression changes. Combinatorial indexing-based single cell technologies allow profiling millions of single cells in each experiment, at substantially reduced cost compared to standard 10X protocols ^24,33,45,46^. To show that our monoclonal screening system in principle enables profiling many EBs at scale and is compatible with combinatorial indexing-based single cell technologies without direct gRNA capture, we shallowly profiled both experiments (**Fig. 4a**) using three-level sci-RNA-seq^45,46^ and enriched for the organoid BC, yielding 46,754 cells (**Fig. 4j**) (hereafter referred to as the ‘**9TF-monoclonal-sci**’ experiment; **Table 1**). In the ’arrayed’ experiment, 4,315 cells (17.3%) were assigned to 49 clonotypes with at least 50 cells, averaging 88 ± 58 cells and ranging from 50 to 302 cells per clonotype. In the ’pooled’ experiment, 3,615 cells (16.5%) were assigned to 42 clonotypes with at least 50 cells, averaging 86 ± 41 cells and ranging from 50 to 215 cells per clonotype. Perturbation identities for each clonotype were determined from the organoid BCs. This is possible because as established above, organoid BCs can identify the linked gRNAs without direct capture (which is not as straightforward with sci-RNA-seq). Of note, the depletion of lateral mesoderm and enrichment of cardiomyocytes in *Hand1* gRNA-expressing EBs was reproduced in this experiment (**Fig. 4k**).

Taken together, while the platform is in need of further optimization, these experiments constitute a proof-of-concept for a monoclonal screening platform. We provide solutions to some of the key challenges encountered in the mosaic screens and establish a scalable method that allows us to identify known roles of master regulators in cell fate commitments, even in heterogeneous multicellular systems. However, our experiments also show that EBs, whether unperturbed, mosaically perturbed or homogeneously perturbed, exhibit considerable inter-individual variation with respect to cell type composition. Particularly as such inter-individual variation is likely pervasive in organoid models, our results reinforce the importance of rigorous assessments of power and reproducibility in genetic screens conducted in mammalian stem cell models.

## Discussion

High-content perturbation screens in cell lines have been highly effective, yet extending these approaches to multicellular systems such as mammalian stem cell-derived organoids presents new challenges. Our study highlights key obstacles that arise in these contexts—particularly the impact of heterogeneity on interpretation—and provides solutions, including a strategy to generate monoclonal embryoid bodies (EBs) that are barcoded with respect to both identity and perturbation within pooled cultures. Based on our findings, we propose best practices for future studies and illustrate the potential of this approach for next-generation high-content screening in multicellular models.

### Challenges, solutions, and best practices for multicellular screens

Applying CRISPR-based perturbation screens to mosaic EBs—a common intermediate in many organoid protocols—we encountered three major challenges. The first challenge was <u>lentiviral silencing</u>. Despite using a promoter reported to resist silencing during differentiation^29,30^, our lentiviral constructs were extensively silenced in differentiating EBs, substantially reducing the fraction of cells that could be analyzed. The silencing affected both targeting and non-targeting constructs, indicating it was not a selective adaptation to specific perturbations. An enrichment of less differentiated cell types (*e.g.* epiblast, PGCs) among GFP-positive cells suggests differential silencing across lineages. While continuous drug selection did not prevent silencing, switching to a piggyBac transposon-based system enabled stable long-term expression and eliminated the need for FACS sorting. This aligns with previous reports that transposon-based delivery can mitigate silencing effects in differentiation contexts^48,49^.

The second—and perhaps more critical—challenge was <u>stochastic clonal skewing</u>, wherein individual EBs exhibited highly uneven representation of perturbations. Despite being initially composed of hundreds to thousands of cells, each mosaic EB was overwhelmingly dominated by a handful of clones, confounding the interpretation of perturbation effects. These imbalances were not solely attributable to selective effects, as they were also observed for NTCs. Instead, we speculate that they arise from either skewed clonal expansion within polyclonal aggregates or significant cell death during EB cavitation^50^ or differentiation. Combined with substantial inter-EB heterogeneity in cell type composition, as well as potentially heterogeneous knockdown efficiencies, this stochastic clonal expansion undermined the reproducibility of perturbation effects. While strong perturbations (*e.g. Carm1*, *Trp53*) rose above this noise, it was difficult to distinguish true effects from false positives. Averaging across large numbers of EBs may help mitigate the consequences of stochastic skewing, but is only practical for smaller libraries. Alternative culturing methods may help reduce heterogeneity^51^, but are unlikely to eliminate it entirely. Of course, it remains possible that other organoid models that do not rely on EBs as an intermediates are more robust to this phenomenon.

To address these issues more directly, we developed a strategy to barcode monoclonal EBs within pooled experiments. The monoclonal aspect eliminates “intra-EB” genotype variability. The barcoding aspect enables us to quantify “inter-EB” variability in the context of a given perturbation, or even no perturbation (*i.e.* baseline organoid heterogeneity).

Based on our experiences, we recommend the following best practices for high-content screening in multicellular systems such as stem cell-derived organoids—whether using polyclonal (*i.e.* mosaic) or monoclonal ESC aggregates: 1) Begin with pilot experiments to assess silencing and heterogeneity using NTC-only libraries; 2) Consider transposon-based delivery systems, genomic safe harbor integration and/or continuous selection to reduce silencing; 3) Leverage quantification of clonal heterogeneity in pilot experiments to inform power calculations for the main experiment(s); 4) In the main experiment(s), include robust internal controls, such as a high proportion of NTCs (*e.g. ∼*10% of all guides), multiple guides per target (*e.g.* at least three), and biological replicates (*e.g.* at least two, ideally more).

### Contrasting monoclonal and mosaic organoids for perturbation screens

To date, nearly all pooled sc-RNA-seq screens using EBs or organoid intermediates have been mosaic in design^12–15^, *i.e.* reliant on polyclonal aggregates of ESCs bearing heterogeneous perturbations. Mosaic perturbations can be avoided by starting each organoid from a single cell bearing a single perturbation, but to our knowledge this has only been pursued in systems such as organoids derived from precursors extracted from intestinal tissue, in which the standard protocol yields clonal organoids^52^ and not by recovering organoid identity alongside single cell profiles.

Mosaic screens are straightforward to implement and in principle, enhance power, *e.g.* given a fixed number of EBs or organoids to be profiled—usually the rate-limiting factor due to cost—a larger number of perturbations can be assessed in a mosaic system, as compared with a system in which each EB or organoid intermediate uniformly bears a single perturbation. The two most obvious disadvantages of mosaic screens relate to cell non-autonomous effects: 1) we cannot measure the effects of a given perturbation on cells’ neighbors; and 2) we cannot measure the effects of having the same perturbation throughout the organoid. These limitations constrain the ability of mosaic screens to model monogenic human developmental disorders, wherein disruptions of specific genes are constitutional. Our results demonstrate an additional major disadvantage -- pervasive stochastic clonal skewing -- which upends the presumed advantages of mosaic screens with respect to the cost required to achieve a given power, at least in EBs and EB-derived organoid models.

Importantly, even after resolving issues of intra-organoid clonal skewing through monoclonal EBs, we observed substantial variation in cell type composition across monoclonal EBs bearing the same perturbation, or even in the absence of perturbation. This kind of “inter-individual” variability—variation across replicate organoids with identical perturbations—would have been difficult to detect, let alone quantify, in traditional mosaic systems. By uniquely barcoding each monoclonal EB, we were able to directly observe and begin to characterize this variability, which appears to be an inherent feature of some organoid models rather than a technical artifact, and has previously been noted based on *in silico* modeling as a potential confounder of organoid perturbation screens^53^. In a companion study^54^, we extend the strategy described here to generate monoclonal gastruloids, and leverage DNA Typewriter^55^ to investigate the origins of inter-individual organoid heterogeneity.

Interestingly, a recent study of mosaic, human iPSC-derived brain organoids encountered related challenges, with EB-derived, multi-donor human brain cortical organoids exhibiting extensive skewing^56^. The authors solved this by initiating brain organoid derivation on single-donor-derived stem cell aggregates, but then disaggregating and pooling these several weeks into the protocol. In the ensuing weeks of differentiation, representation of clones derived from multiple donors is well maintained within individual brain organoids, termed “chimeroids”. Taken together with our results, the implication is that stochastic clonal skewing is a phenomenon that occurs early in EB differentiation.

Although monoclonal embryoids have been described previously^57^, they remain underutilized. Our work demonstrates that they offer several key advantages for perturbation screens: 1) <u>The elimination of intra-EB</u> <u>genotype imbalance</u>: All cells within a monoclonal EB carry the same perturbation, removing a major confounder; 2) <u>Dual barcoding</u>: Each EB can be tagged with a high-complexity barcode that encodes both perturbation identity and EB-of-origin, enabling per-EB comparisons across perturbation conditions; 3) <u>Quantification of reduced penetrance and variable expressivity</u>: By tagging monoclonal EBs such that cell type compositions can be compared across many ‘individuals’ exposed to the same culture conditions in the same dish, there is the potential to study these phenomena, which are pervasive in Mendelian disorders^58^. 4) <u>Versatility</u>: Since EBs serve as intermediates for many organoid systems, this platform could be extended to generate barcoded monoclonal organoids of many kinds, useful for disease modeling, drug screening and lineage tracing; and 5) <u>Scalability</u>: Our proof-of-concept demonstrates the potential of barcoded, monoclonal organoids to support thousands of perturbations, particularly when paired with combinatorial indexing technologies for cost-efficient single-cell profiling. This opens the door to comprehensive TF perturbation screens with high coverage across cell types.

### Outlook

Looking ahead, monoclonal barcoded organoids offer a powerful platform for scalable, high-content genetic screening in complex multicellular systems. Yet several aspects of the system will benefit from further optimization. These include improved recovery of organoid barcodes in sci-RNA-seq workflows, more efficient generation of truly monoclonal EBs, and strategies to better manage inter-individual heterogeneity in both cell type composition and perturbation efficacy. Nevertheless, the ability to disentangle cell-intrinsic effects, track per-organoid phenotypes, and scale to thousands of perturbations makes this a promising platform. With continued refinement, barcoded monoclonal organoids could become a foundational tool for dissecting the functional genomics of development. Furthermore, we envision that this approach may enable scaling of iPSC-derived human organoids, facilitating modeling how both common and rare variants shape molecular and cellular phenotypes in health and disease.

## Data & code availability

The data generated in this study can be downloaded in raw and processed forms from the NCBI Gene Expression Omnibus under accession number <u>GSE291368</u>. The code used here is available at: https://github.com/shendurelab/EBTF.

## Acknowledgments

We thank the members of the Shendure lab for helpful discussions. This work was supported by the Brotman Baty Institute for Precision Medicine and grants from the Paul G. Allen Frontiers Group (Allen Discovery Center for Cell Lineage Tracing to J.S. and C.T.), Seattle Hub for Synthetic Biology, a collaboration between the Allen Institute, Chan Zuckerberg Initiative and University of Washington (award number CZIF2023-008738 to J.S. and C.T.), European Molecular Biology Organisation (ALTF 732-2017 to S.D.), Human Frontier Science Program (LT 000213/2018 to S.D.), Damon Runyon Cancer Research Foundation (DRG-2435-21 to J.-B.L.), Alex’s Lemonade Stand Foundation (Grant 19-15730), Crazy 8 Initiative (to J.S.), and National Institutes of Health (1UM1HG011586 to J.S.; R01HG010632 to J.S. and C.T.; F31HG011576 to S.G.R.). J.S. is an Investigator of the Howard Hughes Medical Institute.

## Author Contributions

S.D., S.R., J.S. designed the research.

S.R., S.D. performed the experiments, with assistance from B.K.M., J.B.L.

C.Q., S.D., S.R. performed the computational analyses, with assistance from J.B.L, A.K., M.D., C.T. S.D., C.Q., J.S. wrote the manuscript, with assistance from S.R.

S.D., J.S. supervised the project.

## Methods

### Experimental methods

#### Mouse ES cell culture

All ES cell lines were derived from the WD44 mouse ES cell line (male homozygous C57BL/6), which was a gift from Dr. Christine Disteche at the University of Washington. Mouse ES cells were maintained on 10 cm dishes precoated with 0.2% gelatin solution in serum/LIF medium consisting of DMEM (Thermo Fisher, cat#10313021), 15% fetal bovine serum (Biowest, Premium bovine serum, S1620), 1% non-essential amino acids (Thermo Fisher, cat#11140050), 1% GlutaMAX (Thermo Fisher, cat#35050061), 0.01% leukemia inhibitory factor (Sigma-Aldrich, ESGRO Recombinant Mouse LIF Protein ESG1107), and 0.001% β-mercaptoethanol (Thermo Fisher, cat#21985023). Cells were passaged at 60-70% cell confluence or every second day.

#### Mosaic embryoid body cell culture

WD44 ES cells were washed twice with PBS (without Mg^+2^, Ca^+2^, -/-) and dissociated into a single-cell suspension using 0.05% trypsin, followed by trypsin inactivation with serum/LIF medium. The cells were then centrifuged at 300g for 5 minutes and resuspended in cell aggregation (CA) medium (no LIF), consisting of DMEM supplemented with 10% fetal bovine serum, 1% GlutaMAX, 1% NEAA, and 0.001% β-mercaptoethanol. Cells were plated at a starting density of 3x10^6^ cells per plate in CA medium on non-adherent 10 cm dishes (Greiner Bio-One, cat#633180). During the first 24 hours, cells self-aggregated into three-dimensional embryoid bodies (EBs). The EBs were maintained in CA medium and passaged every two days. For passaging, EBs were carefully collected using a 25 mL serological pipette and transferred to a 50 mL tube. The EBs were allowed to settle at the bottom of the tube by gravity, after which the old medium was gently removed. Fresh CA medium (12 mL) was then added to the tube, and the EBs were transferred back to a new culture dish. By day 14, EBs exhibited cystic structures, a morphological change consistent with differentiation.

### Lentiviral transduction

Lentiviral transductions were performed similarly across all experiments. One day prior to lentiviral transduction, mESCs were seeded on a 6-well plate at a density of 400,000 cells per well and cultured in serum/LIF medium. On the following day, old medium was removed and replaced with fresh medium with 0.1% protamine sulfate (8 mg/ml, Sigma P-45605) used as an adjuvant. High titer virus, produced by the Fred Hutchinson Vector Lab, was added to each well at increasing amounts to achieve the desired multiplicity of infection (MOI). After adding virus, cells were again transduced 8-10 hours later with the same amount of virus and protamine sulfate added directly to the medium. After 24 hours from the initial transduction, old medium was exchanged and cells were maintained and passaged at 48 hours post transduction. Cells were FACS (via BD FACSAria II, gating on GFP) analyzed to estimate MOI for different virus concentrations prior to continued maintenance with or without selection (see Methods section ‘Perturbation screens in mosaic EBs’ below).

### Making monoclonal cell lines

WD44 was the parental cell line used for generating monoclonal CRISPRcut and CRISPRi lines. Monoclonal lines were generated by transgene integration using transduction with lentivirus, produced by the Fred Hutchinson Vector Lab, and prepared from plasmid constructs comprising CRISPRcut (Cas9 and blastomycin resistance gene, Addgene# 52962) or CRISPRi^59^ (dCas9-BFP-KRAB fusion, Addgene# 46911) machinery. For CRISPRcut, selection was performed using blastomycin (Thermo Fisher, A1113903) at 5ug/ml, whereas for CRISPRi, no antibiotic selection was applied. Following transduction, CRISPRcut cells underwent individual colony picking and expansion in separate wells of a 12-well plate. Stable construct expression was evaluated by qPCR for the Cas9 gene, and the clone with the highest expression was selected for downstream experiments. In contrast, CRISPRi cells were initially grown polyclonally, sorted based on BFP fluorescence via flow cytometry (BD FACSAria II), and then enriched through a second sorting step before sparse plating for colony picking. FACS analysis was performed on each monoclonal line to confirm strong, stable and unimodal BFP expression.

### Cloning and transduction of CROP-seq gRNA libraries

We used the CROP-seq vector^28^ to deliver and stably integrate gRNAs into cells, with which the gRNA also becomes part of the puromycin-resistance mRNA transcribed by RNA polymerase II and thus detectable in standard scRNA-seq protocols. To estimate transduction efficiency and determine the MOI in our mosaic EB perturbation screens, we modified the CROP-seq vector by incorporating a p2A-GFP sequence at the 5′ end of the puromycin cassette. For gRNA design, we relied on two primary databases: for CRISPRcut, we utilized the Brie database^60^ to select highly active gRNAs with minimized off-target effects. For CRISPRi screens, we used the Horlbeck database^61^, selecting the top three scoring gRNAs per gene. For gene targets not covered in the Brie or Horlbeck databases, we supplemented with the Broad GPP database (https://portals.broadinstitute.org/gpp/public/).

The ‘pilot-mosaic’ screen targeted 31 TFs important for ES cell maintenance or EB formation/development (**Supplementary Table 3**), with three gRNAs per TF for both CRISPRi and CRISPRcut, resulting in a total of 93 targeting gRNAs as well as 51 non-targeting controls (NTCs) (Different NTC gRNAs were used for CRISPRi and CRISPRcut; **Supplementary Table 3**). For the ‘125TF-mosaic’ screen (CRISPRcut only), 125 TF targets were included with three gRNAs per TF, yielding 375 targeting gRNAs and 81 NTCs (**Supplementary Table 3**). The ‘allTF-mosaic’ screen targeted 1,644 TFs with three gRNAs per TF, resulting in 4,932 targeting gRNAs and 999 NTCs (**Supplementary Table 3**). Note that targeting and NTC ratios were adjusted in cloning to reflect a 80:20 ratio (see below). For an overview on the experimental design of all screens, see **Table 1**.

For the ‘pilot-mosaic’ and ‘125TF-mosaic’ screens, gRNAs were ordered as standard desalted single-stranded oligos from IDT, pooled to a concentration of 1 ng/μL, and amplified using Kapa Biosystems HiFi Hotstart ReadyMix (KHF) with the following primers: Forward (5′-ATCTTGTGGAAAGGACGAAACA) and Reverse (5′-CTGTTTCCAGCATAGCTCTTAAAC) at an annealing temperature of 63.5°C. The amplified products were purified using Zymo Research DNA Clean & Concentrator-5 kits. For the ‘allTF-mosaic’ screen, gRNAs were ordered as a pool from CustomArray, where each unique oligo was duplicated six times, resulting in a total of 91,290 oligos. This pool included both CRISPRi and CRISPRcut oligos and was designed to allow amplification of subpools for targeting gRNAs or non-targeting controls using dial-out retrieval primers. Subpools were purified with Zymo Research DNA Clean & Concentrator-5 kits and further amplified using inner primers identical to those used in the other screens with 1 ng as input. For all screens, separate pools of targeting guides and NTC were mixed at a ratio of 80:20 prior to cloning into the modified CROP-seq vector.

The modified CROP-seq vector was prepared for Gibson assembly by digestion with BsmBI and alkaline phosphatase (FastDigest Esp3I and FastAP, Thermo Fisher), followed by gel extraction to remove filler sequences and purification using Zymo Research DNA Clean & Concentrator-5 kits. Purified gRNA amplicons were assembled into the digested CROP-seq vector using the NEBuilder HiFi DNA Assembly Cloning Kit (NEB) at a molar ratio of 100 fmol vector:200 fmol insert. Gibson reactions were incubated at 50°C for 15 minutes. Assembled products were transformed into Stable Competent E. coli (NEB C3040H), with sufficient coverage to ensure >20 transformant clones per gRNA in the library. Transformed colonies were cultured, and the plasmid DNA library was prepared using ZymoPURE Maxiprep kits, followed by final purification with Zymo Research DNA Clean & Concentrator-5 kits for lentiviral production and transduction (see Methods section ’Lentiviral transduction’ above).

### Perturbation screens in mosaic EBs

For all mosaic screens, we applied a largely identical perturbation workflow and highlight differences between screens here and in **Table 1**. Briefly, we stably integrated gRNAs into mESC monoclonal CRISPRi or CRISPRcut cell lines using lentivirus produced from libraries of modified CROP-seq plasmid DNA containing pooled gRNAs (*e.g.* gRNAs for pilot-mosaic, allTF-mosaic, or 125TF-mosaic screens). Using the GFP reporter on the modified CROP-seq construct, we could estimate transduction efficiency via FACS analysis, wherein we titrated viral titers such that infectivity of mESCs yielded a cell population of <10% GFP positivity. Based on an idealized Poisson distribution, we estimate that ∼95% of cells that are GFP positive express at most 1 gRNA. For the pilot screen, the TF targets chosen are literature validated essential genes (e.g. for pluripotency maintenance, EB formation, and germ layer differentiation) and therefore we chose not to enrich the GFP-positive mESC population with puromycin selection to avoid markedly reducing EB formation and growth dynamics. However, for the allTF-mosaic and 125TF-mosaic screens, we intuited that most perturbed TF targets would not have a significant impact on EB formation or growth. For the allTF-mosaic and 125TF-mosaic screen, we enriched the population of successfully transduced cells (as measured by GFP fluorescence) with 2ug/mL puromycin selection for 5 days, reaching ∼100% GFP positive rate. Cell pellets were collected after stable integration and/or puromycin selection for PCR amplification of gRNAs from genomic DNA to determine gRNA representation in the cell population upon EB induction. After confirming an even representation of gRNAs in the cell population, mESCs were differentiated to EBs as described above. At day 21, EBs were dissociated into single-cell cells and processed following the 10X Genomics protocol.

### Single cell preparation for 10X Chromium

To obtain a single-cell suspension for downstream single-cell experiments, EBs at a given time point, typically at day 21 for the mosaic perturbation screens (pilot-mosaic, allTF-mosaic, and 125TF-mosaic) or at day 7, 14, or 21 for the wildtype time series experiment, were collected into 50 mL conical tubes at a density of three plates per tube and allowed to settle by gravity. The old CA medium was carefully removed and replaced with PBS (-/-) for two sequential washes, allowing EBs to settle by gravity after each wash. After removing the final wash, 3 mL of 0.25% trypsin was added to the tube, and the EBs were incubated at 37C for 5 min with gentle agitation. To achieve a uniform single-cell suspension, the trypsinized EBs were carefully pipetted up and down using a P1000 pipette followed by quenching with CA medium. The cell suspension was passed through a 40 um cell strainer, centrifuged at 300g for 5 min, and the cell pellet resuspended to a final concentration of 2 million cells / mL. FAC sorting was performed to enrich for a clean GFP positive single-cell population, after which the cells were centrifuged at 400g for 5 min and resuspended in cold PBS containing 0.04% BSA at a concentration of approximately 1200 cells/uL. The resulting single-cell suspension was then processed following the 10X Genomics protocol as per the manufacturer’s instructions, loading 16K cells per lane for an estimated recovery of 10K cells per 10X lane. Sequencing was performed as described in **Table 1**.

### gRNA enrichment in cDNA libraries via hemi-nested PCR

To enrich gRNA sequences from the 3′ UTR of puromycin resistance gene transcripts generated by the CROP-seq integrant, a three-step hemi-nested PCR protocol was employed^62^. Amplification was carefully monitored in real time using qPCR with SYBR Green (Invitrogen) to prevent overamplification, and reactions were halted prior to reaching saturation.

#### Step 1: Initial PCR

Approximately 10–13 ng of full-length 10X scRNA-seq cDNA was used as input for the first PCR reaction, carried out in a 50 µL volume with KHF polymerase (annealing temperature: 65°C). To monitor amplification, SYBR Green was added to the reaction mix (Forward Primer: 5′-TTTCCCATGATTCCTTCATATTTGC-3′, Reverse Primer: 5′-ACACTCTTTCCCTACACGACG-3′).

#### Step 2: Intermediate PCR

The initial PCR was purified using Agencourt AMPure XP beads (1x ratio, Beckman Coulter). A 1/25th aliquot of the purified pooled product served as input for the second PCR step. This reaction was conducted in a 50 µL volume using KHF polymerase and included SYBR Green for qPCR monitoring. Amplification was performed at an annealing temperature of 65°C (Forward Primer with P7 Adapter: 5′-GTCTCGTGGGCTCGGAGATGTGTATAAGAGACAGcTTGTGGAAAGGACGAAACAC-3′, Reverse Primer (R1-P5): 5′-AATGATACGGCGACCACCGAGATCTACACTCTTTCCCTACACGACG-3′).

#### Step 3: Final PCR

The products of the second PCR were purified again using a 1x AMPure XP bead clean-up, and 1/25th of this pooled product was used as input for the final PCR reaction. This step was performed in a 50 µL volume with KHF polymerase and SYBR Green for qPCR monitoring, with an increased annealing temperature of 72°C (Forward Primer (Nextera P7 Indexing Primer): 5′-CAAGCAGAAGACGGCATACGAGAT(index)GTCTCGTGGGCTCGG-3′, Reverse Primer (R1-P5): 5′-AATGATACGGCGACCACCGAGATCTACACTCTTTCCCTACACGACG-3′). The final product was purified using 1x AMPure XP beads.

### Quantifying gRNA representation in individually picked mosaic EBs

To assess bottlenecking in mosaic EBs, we analyzed gRNA representation in day 21 EBs harboring the 125TF gRNA library. Briefly, mESCs transduced with the 125TF library were differentiated into EBs (see Methods section ‘Mosaic embryoid body cell culture’ below). On day 21, we randomly hand-picked EBs with a pipet tip under a microscope, counted total cell numbers per EB after trypsinization using a hemocytometer and trypan blue staining, and separately collected 12 additional EBs into individual PCR tubes. Genomic DNA (gDNA) was extracted using the PicoPure DNA Extraction Kit (Applied Biosystems), followed by PCR amplification (Forward = GTCTCGTGGGCTCGGAGATGTGTATAAGAGACAGTGTGGAAAGGACGAAACACC, Reverse = TCGTCGGCAGCGTCAGATGTGTATAAGAGACAG(NNNNNNNNNN)CGGTGCCACTTTTTCAAGTT, Tm = 65, Enzyme = KHF) and real-time quantification using qPCR with SYBR Green (Invitrogen). We included a UMI in the reverse primer to assist with deduplication of PCR products. As a reference, we also extracted gDNA from the initial mESC population transduced with the gRNA library and PCR amplified gRNA similarly as above. A standard curve was generated by collecting mESCs in increasing amounts (100, 1K, 5K, 10K, 50K, 100K cells) to assess our ability to quantify gRNA read frequency at different cell numbers. The resulting amplicons were further amplified using Nextera primers and sequenced on a NextSeq 500 to quantify gRNA representation across samples.

### Quantification of gRNAs from genomic DNA

For all screens, stably transduced mESCs, along with their corresponding EB samples, were collected as cell pellets. Genomic DNA was extracted from these pellets using the Qiagen DNeasy Blood and Tissue Kit, following the manufacturer’s protocol. The isolated genomic DNA served as the template for a nested PCR amplification of the integrated gRNA sequences, performed in two sequential rounds to ensure specificity and optimal library preparation. Guides were also amplified from the plasmid library. All PCR reactions were performed using Kapa HiFi 2X Master Mix and closely monitored as qPCR reactions by spiking with SYBR Green (Invitrogen) to prevent overamplification. Amplifications were halted before saturation to maintain library integrity. To confirm the success of amplification, 1 µL of each PCR product was analyzed on a 6% TBE polyacrylamide gel (Invitrogen). In the first PCR round, 200 ng of genomic DNA or 10 ng of plasmid DNA was used as input (Forward Primer: 5′-GTCTCGTGGGCTCGGAGATGTGTATAAGAGACAGTGTGGAAAGGACGAAACACC-3′, Reverse Primer: 5′-TCGTCGGCAGCGTCAGATGTGTATAAGAGACAGCGGTGCCACTTTTTCAAGTT-3′). After the first PCR, amplified products were purified using 2x AMPure XP beads (Beckman Coulter) to remove residual primers and nonspecific products. The purified PCR products were then used as templates for the second round of PCR, with 1/25th of the cleaned product serving as input. This second PCR step employed Nextera P7 and P5 primers. As in the first round, amplification was done with Kapa HiFi 2X Master Mix along with SYBR Green to monitor amplification via qPCR, ensuring precise endpoint determination. The product was cleaned up with Zymo Research DNA Clean & Concentrator-5 and sequenced on a Nextseq 500 75-cycle high-output kit (R1:30 bp, I1:8 bp, I2:8 bp, R2:41 bp).

### PiggyBac transfection

PiggyBac transfections were performed using a reverse transfection protocol with Lipofectamine 2000 Reagent (Invitrogen). Briefly, mESCs cultured on 10 cm dishes were washed twice with PBS (-/-), treated with 2.5 mL of 0.05% trypsin, and incubated at 37°C for 5 minutes. Cells were then pipetted up and down to ensure single-cell dissociation, after which trypsin was inactivated with 7.5 mL of serum/LIF medium. The cell suspension was strained through a 40 μm filter, centrifuged at 300g for 5 minutes, and resuspended in serum/LIF medium at a final density of 500,000 cells/mL. In parallel, transfection reagents were prepared by mixing 10 μL Lipofectamine with 240 μL Opti-MEM and separately preparing a DNA mixture containing piggyFlex and piggyBac transposase at a validated ratio of 0.03:1, optimized to achieve a low multiplicity of integration (∼1 construct per cell). Both mixtures were incubated separately for 5 minutes before being combined and incubated for an additional 20 minutes. Transfections were conducted in gelatin-coated 6-well plates. The transfection mixture (500 μL) was first added to each well, followed by 2 mL of mESCs (1M cells) to reach a final volume of 2.5 mL. The plate was gently swirled or lightly tapped to ensure thorough mixing.

### PiggyFlex gRNA libraries

#### piggyFlex

The ’piggyFlex’ construct is a piggyBac-based transposon (Addgene #218234) that comprises a gRNA paired to a complex organoid barcode (organoid BC). The gRNA expression is driven by a Pol3 promoter (hU6) while the expression of the organoid BC (see below, *organoid BC design*), located at the 3’ end of GFP in the puro-p2A-GFP cassette, is driven by a Pol2 promoter (EF1alpha) located upstream the cassette. The gRNA and organoid BC are independently captured alongside single cell transcriptomes using the 10X Genomics capture sequences (CS) in the scaffold (CS1) or at the 3’ end of GFP (CS2), respectively. The complete transposon, 5’-U6-gRNA-EF1alpha-puro-p2A-GFP-organoid BC-3’, is flanked by ITRs and can be stably expressed following genomic integration with piggyBac transposase.

#### piggyFlex functional validation test

Before implementing the piggyFlex construct, we aimed to validate its perturbation efficacy in CRISPRcut mESCs. For this purpose, we selected essential gene targets (Atf1, Baz1b, Brca1, Ccnt1, Cdk7, Hmgb2, Kdm4a, Ncoa3, Nelfcd, Mcm3, Mcm7, Polr2b, Polr2f, Polr2g, Polr2h, Psmc5, Rfc1, Taf13, Taf1b, Uhrf2) that had exhibited the highest fold change depletion in ESCs integrated with the allTF-mosaic screening library. A total of 60 gRNAs (3 per gene target) and 60 NTCs were cloned into the piggyFlex construct. The library was transfected into CRISPRcut mESCs under low MOI conditions to ensure single-copy integrations. Cells were maintained for 14 days to allow stable transposon integration and CRISPR activity.

Cell pellets were collected from two biological replicates, and genomic DNA was extracted using the Qiagen DNeasy Blood and Tissue Kit. gRNAs were amplified from genomic DNA using initial PCR primers designed specifically for the piggyFlex construct (Forward Primer: 5′-GTCTCGTGGGCTCGGAGATGTGTATAAGAGACAGTGTGGAAAGGACGAAACACC-3′, Reverse Primer: 5′-TCGTCGGCAGCGTCAGATGTGTATAAGAGACAGTTGCTAGGACCGGCCTTAAAGC-3′).

Amplifications followed the same conditions as described in the ’Quantification of gRNAs from Genomic DNA’ Methods section. Sequencing was performed on an NextSeq 500 using a 75-cycle high-output kit with the following read structure: R1 (42 bp), I1 (10 bp), I2 (0 bp), and R2 (42 bp). Mapping and creation of read count tables were conducted using QuasR^63^, normalizing each biological replicate to its corresponding plasmid library. This approach ensured accurate quantification of gRNA representation and perturbation efficacy.

### organoid BC design

The organoid BC was designed using two distinct approaches. In the initial design (‘pilot-monoclonal’) (**Table 1**; **Fig. 3a**), the gRNA is paired to an organoid BC composed of a 15N randomer. The pairing is determined by sequencing the plasmid library at an intermediate cloning step to establish the link between the gRNA and the organoid BC. In contrast, the later design (used in ‘9TF-monoclonal-arrayed’, ‘8TF-monoclonal-pooled’, ‘9TF-monoclonal-sci’) (**Table 1**; **Fig. 4**) involved a semi-random barcode in which the first part (8N) of the barcode is synthesized predefined to unequivocally identify the linked gRNA, while the second part (9N) is random and therefore sufficiently complex to serve as an organoid identifier. The latter design eliminates the need for sequencing the plasmid library to determine gRNA-barcode pairings, as the linkage is predefined.

### PiggyFlex cloning for pilot-monoclonal (organoid BC v1)

To construct the piggyFlex vector for monoclonal EB pilot experiments, the piggyFlex plasmid backbone was digested with SalI and BbsI (NEB) in 10X NEBuffer r2 at 37C overnight to ensure complete digestion. This enzymatic digestion separated the vector into two products: the EF1alpha-puro-p2A-GFP cassette and the linear plasmid backbone. The digested products were resolved on a 1% agarose gel in TAE buffer. The linear backbone was gel extracted using a gel extraction kit (NEB), while the EF1alpha-puro-p2A-GFP cassette was reserved for reintegration in a later cloning step.

To prepare the gRNA library for incorporation, the NTC gRNA sequences from the all TF screen were PCR amplified using primers designed to introduce homology arms for Gibson assembly, along with an organoid BC sequence and a SapI site for subsequent scaffold reassembly:

● Forward Primer: 5′-ATCTTGTGGAAAGGACGAAACACCG-3′
● Reverse Primer:

5′-GCCTTAGCCGCTAATAGGTGAGCTTGTCGANNNNNNNNNNNNNNNTCGACTTAACGCGTTCACTTGTACAGCTCGGAAGAGCGCTCCTGCTCTTCTTGCTATGCTGTTTCCAGC-3′

Amplification was performed using Q5 High-Fidelity Polymerase (NEB), and the resulting products were gel purified. The amplified gRNA library was then assembled into the linear backbone using Gibson assembly (NEBuilder HiFi DNA Assembly, NEB). The assembled constructs were transformed into Stable Competent E. coli (NEB C3040H) and plasmid DNA was extracted using a midiprep kit (Zymo Research). At this step, the library was sequenced by NGS to determine gRNA and organoid BC pairings to be incorporated as a lookup table.

The partially assembled piggyFlex vector was digested with SapI for 1 hour at 37C, and gel-purified. In the final assembly step, the purified digestion product was combined with the reserved EF1alpha-puro-p2A-GFP cassette from the gRNA cloning step. This cassette included the 3′ end of the gRNA scaffold and capture sequence 1, and its integration completed the full gRNA scaffold. In the final assembled product, the gRNA is positioned upstream of the EF1alpha-puro-p2A-GFP cassette, and capture sequence 2, encoded in the vector backbone, is located at the most 3′ end, immediately downstream of the organoid barcode (BC). This reaction reconstituted the complete piggyFlex vector (5′-U6-gRNA-EF1alpha-puro-p2A-GFP-organoid BC-3′). The final product was cleaned with AMPure beads, transformed into electrocompetent cells (NEB C3020K), and plasmid DNA was extracted using a midiprep kit. The resulting plasmid library was PCR amplified and sequenced to confirm successful cloning.

*PiggyFlex cloning: ‘9TF-monoclonal-arrayed’, ‘8TF-monoclonal-pooled’, ‘9TF-monoclonal-sci’ (organoid BC v2)* For the piggyFlex experiments targeting transcription factors (TFs), we designed a cloning strategy incorporating nine TF targets (Carm1, Gata6, Hand1, Hes5, Hhex, Lhx1, Lmo2, Six3, and T) and NTCs. Each target and NTC was represented by three distinct gRNAs, totaling 30 unique gRNAs. Individual primer pairs were designed for each TF target and NTC to facilitate precise amplification and cloning into the piggyFlex vector. The forward primers contained the respective gRNA sequence within a standardized scaffold (*5′-ACGAAAGAAGACCTCACC*\*\***GUIDE**\*\*\**GTTTAAGAGCTATGCTGGAAAC-3′*). The reverse primers incorporated a semi-random barcode comprising an 8N predefined sequence uniquely identifying the paired gRNA, followed by a 9N randomer (*5′-GCTCGACATGTTCACTTGCGCAATTCAGCTA8barcodeNNNNNNNNNCAGCTGTTCGA-3′*), to enable clonal EB identification. Amplified gRNA constructs were cloned into the piggyFlex backbone via Gibson assembly. All subsequent cloning steps, including SapI digestion, cassette integration, and final assembly cleanup, were performed as described above for piggyFlex cloning with organoid BC v1. The resulting plasmid library enabled construction of a targeted TF screening library with predefined gRNA-barcode pairings suitable for high-resolution downstream functional analyses.

### Summary of experimental transfections

The experimental design included two biological replicates for the piggyFlex validation experiment and two biological replicates for the pilot-monoclonal experiment. 9TF-monoclonal-arrayed included nine TFs and one non-targeting control, for a total of 10 individual transfection conditions. The 8TF-monoclonal-pooled consisted of a pooled version of the same guides, minus Carm1 targeting guides, due to our concern that these would skew equal gRNA representation. The 9TF-monoclonal-sci experiment consisted of cells from both the above experiments..

### Making monoclonal EBs

Prior to monoclonal EB formation, piggFlex-transfected mESCs were initially cultured as a pool on gelatin-coated 10 cm plates and maintained in serum/LIF medium. Similar to the standard passaging protocol for mESCs, old medium was removed, cells were washed twice with PBS (-/-), and dissociated using 0.05% trypsin. The cell suspension was then passed through a 40 μm filter, centrifuged, and resuspended in fresh serum/LIF medium at a density of 2 million cells/mL. To ensure a single-cell suspension, cells were sorted via FACS into 15 mL tubes and plated at a density of 4,000 cells per well of a 6-well plate for the pilot-monoclonal experiment and 400 cells per well for the 9TF-monoclonal-arrayed and 8TF-monoclonal-pooled experiments. The 6-well plates were pre-seeded 24 hours in advance with inactivated mouse embryonic fibroblasts (MEFs) (Thermo Fisher, cat#A34959) at a density of 400,000 inactivated MEFs per well. Following single-cell plating of the mESCs, gentle shaking was performed to ensure even distribution of mESCs across the wells. Fluorescence microscopy was used to confirm that sorted single cells were GFP-positive and evenly dispersed across the wells. mESCs were maintained in serum/LIF medium for 4–5 days with daily medium changes, during which single cells proliferated into colonies.

To harvest the colonies, the medium was removed, cells were washed twice with PBS (-/-), and 2 mL of collagenase type IV (Stemcell Technologies) was added to each well, followed by incubation at 37°C for 20 minutes. After incubation, gentle agitation by tapping the sides of the plate facilitated colony detachment from the MEF layer. To inactivate collagenase, 2 mL of warm CA medium was added per well, and colonies were transferred into a 50 mL tube using a 10 mL pipette. Colonies were allowed to settle by gravity for 10 minutes, followed by a single wash with CA medium. After resuspension in 30 mL of CA medium, colonies were evenly distributed across two 10 cm low-adherent dishes. Over the next 24 hours, the colonies transitioned from a disk-like morphology to spheroids.

### piggyFlex screens in monoclonal EBs

#### pilot-monoclonal

Two independent replicate transfections were performed and kept separate throughout the experiment. Cells were transfected under low multiplicity of integration conditions, as established in previous PiggyBac optimization experiments, using the piggyFlex construct and a organoid barcode v1 library. This library comprised 5,340 unique NTC gRNAs, each paired with a median of 126 distinct 15N barcodes. Following transfection, monoclonal EBs were generated as described above. From each replicate, 15 monoclonal EBs were hand-picked randomly on day 8, dissociated into single-cell suspensions as described below, and processed separately. Each replicate was loaded onto two 10X Genomics lanes.

#### 9TF-monoclonal-arrayed and 8TF-monoclonal-pooled

For the 9TF-monoclonal-arrayed screen in monoclonal EBs, we handpicked 10 EBs per condition (10 conditions, 9 TFs + NTCs). EBs were chosen based on the following criteria: medium-to-large size (>500 µm), spherical morphology (avoiding irregular or merged EBs), GFP positivity, and visible contraction indicative of cardiac differentiation. For the 8TF-monoclonal-pooled experiment approximately 150 EBs were collected in 3–4 rounds, with approximately 50 EBs picked per round using aspiration with a P1000 pipette. EBs were monitored for GFP expression during selection to ensure optimal quality.

EBs were collected into 2 mL Eppendorf tubes containing CA media, with 5 EBs per tube for conditions 1–10 of the arrayed experiment and 75 EBs per tube for pooled experiment. In total, four tubes were prepared: two containing 50 EBs each (conditions 1–10, arrayed) and two containing 75 EBs each (pooled). These were subsequently combined into two 50 mL conical tubes: one for conditions 1–10 of the arrayed and another for the pooled experiment.

For dissociation into a single-cell suspension, CA media was removed from each 2 mL tube, and 750 µL of 0.25% trypsin was added. Tubes were incubated at 37°C on a thermomixer set to 1100 rpm, with dissociation monitored under an epifluorescence microscope at 5-minute intervals. At the 10-minute mark, gentle trituration with a P1000 pipette was performed to aid dissociation, with additional trypsin added if necessary. Complete dissociation was typically achieved within 20 minutes. Trypsin was inactivated by adding CA media, and the cell suspension was further diluted before being passed through a 45 µm strainer into a 50 mL conical tube containing 10 mL of CA media.

Cells were counted using a hemocytometer, transferred to a 15 mL conical tube for ease of handling, and centrifuged at 400g for 5 minutes. The supernatant was removed, and the cell pellet was resuspended in PBS + 0.04% BSA. The initial cell concentration was adjusted to 2400 cells/µL. A second cell count was performed, and the suspension was further diluted to a final concentration of 1200 cells/µL. Single-cell suspensions from each condition were then processed using the 10X Genomics High Throughput (HT) kit, with cells loaded across 16 lanes. The 10X Genomics direct capture (‘Feature Barcoding’) protocol was used to detect gRNAs. For the pilot-monoclonal, 9TF-monoclonal-arrayed, and 8TF-monoclonal-pooled experiments, organoid BC sequences were additionally enriched from the cDNA as described in the section below. Final libraries were sequenced together on a NextSeq2000 (**Table 1**).

### Organoid barcode enrichment in cDNA libraries via nested PCR

Organoid BC sequences were enriched from piggyFlex-derived transcripts captured in poly(dT)-primed 10x scRNA-seq cDNA, using a three-step nested PCR protocol analogous to the approach described in Methods section ‘gRNA enrichment in cDNA libraries via hemi-nested PCR’ above. As with the CROP-seq construct, where gRNAs are transcribed and recovered from the 3′ UTR of a reporter gene, the organoid BC is transcribed as part of the piggyFlex transcript and recovered from the resulting cDNA. An initial PCR was performed using primers oJBL207 (5′-CTACACGACGCTCTTCCGATCT-3′) and oSR38 (5′-GTGAACCGCATCGAGCTGAA-3′), followed by a semi-nested PCR with oJBL324 (5′-AATGATACGGCGACCACCGAGATCTACACTCTTTCCCTACACGACGCTCTTCCGATCT-3′) and oJBL529 (5′-ATCCATGGCCCTCAGGCATACTGTTCCAACTCCAGGGCACAAGCTGGAGTACAAC-3′). A final indexing PCR was performed using oJBL76 (P5) (5’-AATGATACGGCGACCACCGAGATCTACAC-3’) and custom-indexed P7 primers (5′-CAAGCAGAAGACGGCATACGAGAT(10 bp index)ATCCATGGCCCTCAGGCATAC-3′) for each sample. All reactions were carried out with KAPA2G Robust HotStart ReadyMix in the presence of SYBR Green for real-time qPCR monitoring, and reactions were halted before the inflection point to prevent overamplification. AMPure beads were used to purify products between each step, and final libraries were used for sequencing. For sequencing, libraries were run with the following configuration: read 1 used the standard Illumina TruSeq primer (≥28 cycles) to read the cell barcode and UMI, index 1 was read with a custom primer (oJBL534, 5′-tggagttggaacaGTATGCCTGAGGGCCATGGAT-3′) for 6–10 cycles to capture the sample index, and Read 2 used a custom primer (oJBL334) for ≥15 cycles to read the organoid barcode.

### 9TF-monoclonal-sci

The sci-RNA-seq3 protocol was performed as previously described^46^, with minor modifications detailed below. First, we used fixed whole cells instead of nuclei, incorporating additional pre-processing steps before the sci-RNA-seq3 experiment. Sample preparation and fixation proceeded separately for the hand-picked monoclonal EBs from the 9TF-monoclonal-arrayed experiments (separately grown conditions 1-10, pooled upon collection) and the 8TF-monoclonal-pooled experiment.

Following trypsin treatment to get single-cell suspension (one 5 mL PBS wash, 750 uL trypsin 0.25% for 5 min at 1200 rpm at 37C on thermal mixer, followed by P1000 trituration, trypsin quenching with pre-warmed 8 mL EB medium, straining through 40 um filter), whole cells were spun down at 4C for 3 min at 500 g, washed with PBS, spun down again and resuspended in 1 mL ice cold PBS + DEPC (10 uL/mL). Cells were then fixed/permeabilized by adding 4 mL methanol + DSP (100 uL of 50 mg/mL DSP in 4 mL methanol), and kept on ice with occasional swirling. Cells were then rehydrated with 2 volumes of SPBSTM. We saw some large DSP crystals in the samples upon fixation (as is common with DSP). Crystals were efficiently removed by (1) gravity sedimentation and (2) filtering with a 40 um strainer. This thus led to 4 different samples for sci: arrayed_gravity, arrayed_filter, pooled_gravity, pooled_filter. Cells were spun down, and resuspended in 1mL SPBSTM, counted, spun down, the medium aspirated and snapped frozen on liquid nitrogen for cold storage.

To perform sci-RNA-seq3, cells were thawed on ice, resuspended in SPBSTM, and the protocol (with whole cells) proceeded as described in Martin et al. 2022^46^. About 1.2M cells total were loaded on the first indexed reverse transcription (RT) plate, with the RT well marking sample identity [arrayed_gravity, arrayed_filter, pooled_gravity, pooled_filter]. To enable enrichment for barcoded constructs, a third sci index (PCR) was also added on the P5 side (Truseq), and primer oJBL315 (5’-CACTCTCGGCATGGACGAG-3’) was spiked in the indexed PCR for reporter enrichment (16 cycles, NEBNext with gap fill step). Following pooling of all wells and 0.8x ampure clean up for final gene expression size selection, a secondary semi-nested PCR with primers P5 and P7 primers oJBL076 (5’-AATGATACGGCGACCACCGAGATCTACAC-3’) +oJBL618-619 for technical PCR duplicates (5’-caagcagaagacggcatacgagatGATCAGggcatggacgagctgtacaagt-3’, 5’-caagcagaagacggcatacgagatTAGCTTggcatggacgagctgtacaagt-3’) was performed with Kapa Robust (12 cycles of amplification, 2 uL of pooled & cleaned up PCR1 samples) to finalize sci-tagged barcoded constructs.

Gene expression and barcode libraries were sequenced together (**Table 1**): gene expression using the same sequencing primers as Martin et al 2022^46^, barcoded libraries using a combination of Illumina primers and custom primers: Read1 (=ligation sci-BC2, UMI, RT sci-BC1) with Truseq Read1; Index1 (=technical replicate index) with oJBL370 (5’-TCGACTTAACGCGTTCACTTGTACAGCTCGTCCAT-3’), Index2 (=PCR sci-BC3) with Truseq index 2; Read2 (=organoid barcode) with oJBL334 (5’-TCGGCATGGACGAGCTGTACAAGTGAACGCGTTAAGTCGA-3’).

### Computational methods

#### Data processing for the wild-type mouse EBs

The single-cell transcriptome data from wild-type mouse EBs at days 7, 14, and 21 were processed using Cell Ranger 7.2.0^64^ with refdata-cellranger-mm10-3.0.0 as the reference. Cells from each day were profiled using an independent lane and processed separately. We extracted the gene-by-cell matrix from the ’raw_feature_bc_matrix’ folder, filtered out cells with UMI counts below 500 or fewer than 250 detected genes, and retained genes from chromosomes 1-19, X, Y, and MT. We then applied the Scrublet/v0.1^65^ pipeline to detect doublets and calculate the percentage of reads mapping to either mitochondrial (i.e. MT%) or ribosomal chromosomes (i.e. Ribo%) for individual cells. After manually examining the distribution of UMIs and MT% across cells, we applied criteria to further remove potentially low-quality cells: those with UMI counts fewer than 2000 or exceeding the top 0.5% of the total cells, doublet scores calculated by Scrublet over 0.2, or MT% over 10% were removed.

After combining cells from days 7, 14, and 21, we applied Seurat/v5^37^ to this final dataset, performing conventional single-cell RNA-seq data processing: 1) retaining protein-coding genes and lincRNA for each cell and removing gene counts mapping to sex chromosomes; 2) normalizing the UMI counts using the SCTransform function^66^ implemented in Seurat; 3) applying PCA and then using the top 30 PCs to calculate a neighborhood graph, followed by louvain clustering; 4) performing UMAP visualization in 2D space (min.dist = 0.3). For cell clustering, we manually adjusted the resolution parameter towards modest overclustering, and then manually merged adjacent clusters if they had a limited number of differentially expressed genes (DEGs) relative to one another or if they both highly expressed the same literature-nominated marker genes (**Supplementary Table 1**). Subsequently, we annotated individual cell clusters using at least two literature-nominated marker genes per cell type label.

#### Integrating and co-embedding cells from mouse EBs, embryos, and gastruloids

Cells from the wild-type mouse EBs and cells or nuclei from two published scRNA-seq datasets of mouse embryos during gastrulation^23^ and early somitogenesis^24^ were integrated using the anchor-based batch correction method in Seurat/v3^26^. The latter dataset originally encompassed the entire organogenesis period, from embryonic day 8 (E8) to postnatal day 0 (P0), but we selected only nuclei from E8 to E10 to align approximately with the developmental stages in the mouse EBs. Additionally, the number of nuclei was randomly downsampled to 200,000 to reduce computational time.

To identify corresponding cell types between mouse EBs and embryos during gastrulation, we first calculated an aggregate expression value for each gene in each cell type by averaging the log-transformed normalized UMI counts of all cells of that type. Next, we applied non-negative least squares (*NNLS*) regression as described previously^45^. Briefly, we predicted gene expression in a target cell type (*T_a_*) in dataset A based on the gene expression of all cell types (*M_b_*) in dataset B: *T_a_ = β_0a_ + β_1a_M_b_*, based on the union of the 1,500 most highly expressed genes and 1,500 most highly specific genes in the target cell type. We then switched the roles of datasets A and B, *i.e.* predicting the gene expression of target cell type (*T_b_*) in dataset B from the gene expression of all cell types (*M_a_*) in dataset A: *T_b_ = β_0b_ + β_1b_M_a_*. Finally, for each cell type *a* in dataset A and each cell type *b* in dataset B, we combined the two correlation coefficients: *β* = 2(*β_ab_* + 0.01)(*β_ba_* + 0.01) to obtain a metric, where high values reflect reciprocal, specific predictivity. We repeated the same analysis to identify correlated cell types between mouse EBs and embryos during early somitogenesis.

Additionally, we also integrated cells from the wild-type mouse EBs and cells from mouse gastruloids from another published dataset^25^ using the anchor-based batch correction method in Seurat/v3^26^.

### Data processing for the ‘pilot-mosaic’ screen dataset

Cells from pooled mouse EBs at day 21 under each of six different conditions were profiled using an independent lane and processed separately: CRISPRcut-unsorted, CRISPRi-unsorted, CRISPRcut-sorted-rep1, CRISPRi-sorted-rep1, CRISPRcut-sorted-rep2, and CRISPRi-sorted-rep2. After standard preprocessing described above, cells with UMI counts fewer than 1000, UMI counts exceeding the top 0.5% of total cells, doublet scores calculated by Scrublet over 0.2, or MT% over 10% for the two unsorted samples and over 20% for the four sorted samples were removed.

Cells from the CRISPRcut-sorted-rep1 sample were removed due to the very limited number of cells recovered. Cells from the remaining five samples were combined with cells from wild-type mouse EBs at day 21, followed by anchor-based batch correction using Seurat/v3^26^. Subsequently, dimensionality reduction using PCA (dims = 30) was performed on the integrated dataset, and UMAP visualization was conducted in 2D space (min.dist = 0.3). Given the relatively small size of this pilot screen dataset, we used a kNN-based approach to automatically transfer cell type labels from previously annotated cells of wild-type mouse EBs instead of *de novo* annotating the cell types. Briefly, cells from the pilot screen dataset were identified with their 10 nearest neighbors in the wild-type sample within PCA space, and each cell was assigned the label of the most abundant cell type among its neighbors.

The sequencing data of gRNAs across cells were processed using Cell Ranger 7.2.0^64^ with default settings. Subsequently, a custom script was employed to extract the read count and UMI count for each gRNA within each cell from the possorted_genome_bam.bam file, with barcode corrections applied using a gRNA whitelist^62^. We manually examined the distribution of UMI counts and read counts across cells, assigning gRNAs to individual cells if their log2-scaled (UMI count + 1) was ≥ 2 and their log2-scaled (read count + 1) was ≥ 5. Of note, to enhance the rigor of the results, only cells assigned with a single gRNA were utilized in the subsequent analysis of the pilot screen dataset.

#### Data processing for the ‘allTF-mosaic’ screen dataset

Cells from pooled mouse EBs, with gRNAs targeting 1643 TFs and NTCs, were profiled using 30 independent lanes of 10X and processed separately. After applying the standard preprocessing described above, cells with UMI counts fewer than 2000, UMI counts exceeding the top 0.5% of total cells, doublet scores calculated by Scrublet over 0.2, MT% over 20% or Ribo% over 40% were excluded.

After combining cells from the 30 10X lanes, we applied Seurat/v5^37^ to this final dataset, performing conventional single-cell RNA-seq data processing: 1) retaining protein-coding genes and lincRNA for each cell and removing gene counts mapping to sex chromosomes; 2) normalizing the UMI counts by the total count per cell followed by log-transformation; 3) selecting the 2,500 most highly variable genes and scaling the expression of each to zero mean and unit variance; 4) applying PCA (dims = 30), followed by utilizing align_cds function implemented in Monocle/v3^45^ with the lane IDs of cells as batch factors to correct for potential batch effects arising from different lanes; 5) using the top 30 corrected PCs to calculate a neighborhood graph, followed by louvain clustering; 6) performing UMAP visualization in 2D space (min.dist = 0.3). Of note, we identified two cell clusters characterized by low UMI counts and high MT%. Moreover, we did not observe highly expressed genes in these clusters corresponding to any known cell types during this developmental stage. Consequently, we removed cells from these two clusters and repeated the aforementioned processing steps. Finally, we annotated individual cell clusters using at least two literature-nominated marker genes per cell type label (**Supplementary Table 1**).

The sequencing data of gRNAs across cells were processed as described above. We manually examined the distribution of UMI counts and read counts across cells, and assigned gRNAs to individual cells based on specific criteria: 1) log2-transformed (UMI count + 1) ≥ 2; 2) log2-transformed (UMI count + 1) < 1.25 × log2-transformed (read count + 1) - 5. In the ’allTF-mosaic’ dataset, we adopted a modified strategy for cells assigned with multiple gRNAs, aiming to preserve as many gRNA assignments as possible: if a cell was assigned both TF gRNAs and NTC gRNAs, the NTC gRNAs were omitted from the assignments. However, if a cell was assigned different TF gRNAs, all those gRNAs were retained. Cells were only counted as NTC cells if they contained single or multiple NTC gRNAs, but no TF gRNAs.

#### Selection of 125 TFs to target in the ‘125TF-mosaic’ experiment

To select the top 125 TFs for which to repeat the experiment in two independent biological replicates (‘125TF-mosaic’ experiment; **Table 1**), we initially applied the strategy detailed below. Of note, we later switched to a different analysis strategy for identifying interesting candidates (see Methods section entitled ‘Identifying TF targets with significant changes compared to NTC’ below), and this is the approach that we present in all of the figures. In the newly implemented approach, we tested 820 targets, including 109 that overlapped with the 125 orginally selected TFs. These overlapping targets had significantly higher neighborhood enrichment scores than the remaining ones (Wilcoxon test, *p* = 5.1 x 10^-31^), implying that the two analysis strategies lead to similar but not identical rankings.

The original strategy is described here for completeness: A Chi-square test, using an expectation maximization approach modeling the functional editing rate of each gRNA^72^, was performed to determine whether the distribution for targeting gRNAs across cell types (clusters) was significantly different compared to NTCs targeting gene deserts. In order to not limit the differential testing to an arbitrary cluster resolution, we also applied the Moran’s I test, a measure of multi-directional and multi-dimensional spatial autocorrelation, to the gRNA x cell matrix. This identifies gRNAs that are significantly spatially correlated compared to NTCs, regardless of cluster identity or resolution. Of note, this spatial correlation is applied in the low-dimensional UMAP space, which is why we later switched to the neighborhood enrichment strategy that identifies neighbors in the high-dimensional PCA space. Nonetheless, the Moran’s I and chi-square q-values correlate (R Spearman = 0.79). For both measures, we compared against the distribution of NTC controls, combining 3 random NTCs at a time to mimic the 3 targeting gRNAs and only selected targeting gRNAs with an empirically determined FDR of < 5%. We further excluded gRNAs whose enrichments across clusters were anticorrelated (R<0.1), to exclude cases where multiple guides for the same target are significantly enriched but in different cell types.

#### Identifying TF targets with significant changes compared to NTC

We employed two strategies to identify targeting gRNAs exhibiting significant changes compared to the NTC. The first strategy utilized cell type annotations to compare the distribution of cells across different cell types between those assigned a specific TF target and the NTC. In the ’allTF-mosaic’ dataset, TF targets (combining three gRNAs) detected in fewer than 50 cells or NTC gRNAs detected in fewer than 20 cells were filtered out. The remaining NTC gRNAs were randomly grouped into sets, each containing three gRNAs, to mimic the three gRNAs for each TF target. Subsequently, cells within each group were randomly downsampled to 100 cells to approximate the cell count for each TF target and to remove biases from differing cell numbers. Chi-squared tests were then conducted to compare the compositions of cell types between cells detected with each TF target and all cells detected with NTC gRNAs (i.e., background). Similar tests were performed to compare cell type compositions between cells detected with each NTC gRNA group and all cells detected with NTC gRNAs. TF targets with p-values below the 0.05 quantile cutoff, as empirically derived from the NTC gRNA group, were considered significant.

The second strategy did not rely on cell type annotations but rather considered the local enrichment of cells to identify substructures within each annotated cell type. In the ’allTF-mosaic’ dataset, TF targets with gRNAs detected in fewer than 50 cells were excluded, and cells associated with each TF target were randomly downsampled to 200 cells. Individual cells were then analyzed to find their 20 nearest neighbors in PCA space (using n = 30 dimensions). For each TF target, the average proportion of nearest neighbor cells sharing the same TF target was calculated.

#### Identifying perturbation-induced gene expression changes

Cells from the lateral plate mesoderm in the ’allTF-mosaic’ dataset were selected, including only those with a single TF target and excluding TF targets with fewer than 50 assigned cells in this cell type. This resulted in 3,696 cells across 13 TF targets (Ctnnb1, Trp53, Cbfb, Lhx8, Nr4a2, Gsc2’, Rfx2, Cecr6, Asz1, Map3k7cl, Hoxa11, Hnrnpr, and Cbx8) and the NTC. Subsequently, Mixscape^38^ analysis was performed using the Seurat/v5^37^ package to identify cells with a targeting gRNA that escaped perturbation and visualize perturbation-specific clusters, employing LDA as the dimensionality reduction method. Differential expression analysis was carried out using the FindMarkers function with min.pct = 0.25 to identify DEGs between cells assigned to each TF target after removal of escaping cells and NTCs. Significant DEGs (avg_log2FC > 0.25 and p_val_adj < 0.001) were further analyzed to determine enrichment in Gene Ontology terms using the topGO package^67^.

#### Data processing for the ‘125TF-mosaic’ screen dataset

Cells from pooled mouse EBs, with 125 TFs screened, were profiled using 32 independent 10X lanes and processed separately. The data were processed using the same strategy as described above, with adjusted criteria to remove potentially low-quality cells after manually examining the distribution of UMIs and other measurements across cells. Specifically, cells with UMI counts fewer than 2000, UMI counts exceeding the top 0.5% of total cells, doublet scores calculated by Scrublet over 0.2, MT% over 20% or Ribo% over 40%.

After combining cells from the 32 10X lanes, we applied Seurat/v5^37^ to this final dataset, performing conventional single-cell RNA-seq data processing as we did for the ’allTF-mosaic’ dataset. Of note, we identified several cell clusters showing quality issues. Moreover, we did not observe highly expressed genes in these clusters corresponding to any known cell types during this developmental stage. Consequently, we removed cells from these clusters and repeated the aforementioned processing steps. Finally, we annotated individual cell clusters using at least two literature-nominated marker genes per cell type label (**Supplementary Table 1**).

The sequencing data of gRNAs across cells were processed using Cell Ranger 7.2.0^64^ with default settings. The subsequent steps for assigning gRNAs to cells were conducted in the same manner as we did for the ’allTF-mosaic’ dataset.

#### Data processing for the ‘pilot-monoclonal’ dataset

The pilot experiment with monoclonal mouse EBs involved profiling 15 manually picked monoclonal EBs. Cells with two technical replicates were sequenced in independent 10X lanes and processed separately. The transcriptome data and gRNA data, including UMI counts for individual features (genes or gRNAs) across cells, were imported into Seurat/v5^37^ as two distinct layers. The gRNAs were filtered out if they were expressed in fewer than 1% of cells, as these likely derive from individual cells floating in the media rather than EBs. The transcriptome data were processed using the same strategy as described above, with adjusted criteria to remove potentially low-quality cells after manually examining the distribution of UMIs and other measurements across cells. Specifically, cells with UMI counts fewer than 1000, UMI counts exceeding the top 0.5% of total cells, doublet scores calculated by Scrublet over 0.2, MT% over 10% or Ribo% over 40%.

After combining cells from the two technical replicates, we applied Seurat/v5^37^ to this final dataset, performing conventional single-cell RNA-seq data processing: 1) retaining protein-coding genes and lincRNA for each cell and removing gene counts mapping to sex chromosomes; 2) normalizing the UMI counts by the total count per cell followed by log-transformation; 3) selecting the 2,500 most highly variable genes and scaling the expression of each to zero mean and unit variance; 4) applying PCA (dims = 30), and then using the 30 PCs to calculate a neighborhood graph, followed by louvain clustering; 6) performing UMAP visualization in 2D space (min.dist = 0.3). Of note, we identified two cell clusters characterized by low UMI counts and high Ribo%. Moreover, we did not observe highly expressed genes in these clusters corresponding to any known cell types during this developmental stage. Consequently, we removed cells from these two clusters and repeated the aforementioned processing steps. Finally, we annotated individual cell clusters using at least two literature-nominated marker genes per cell type label (**Supplementary Table 1**).

### Identifying clonotypes in the ‘pilot-monoclonal’ experiment

In the ‘pilot-monoclonal’ experiment, cells from the two technical replicates were merged. We first examined the joint distribution of log2-scaled UMI counts and log2-scaled read counts for individual organoid barcodes (organoid BCs). Organoid BCs with a log2-scaled read count ≥ 15 were selected (n = 26), and then assigned to individual cells if the UMI count was ≥ 10 and the proportion of UMI counts within each cell was ≥ 30%.

After excluding cells without any assigned organoid BCs (∼13% of cells from replicate-1 were removed), the UMI count across the organoid BCs for individual cells was normalized by the total count per cell, followed by scaling the normalized counts per organoid BC. Pairwise Pearson correlations were then computed for each pair of organoid BCs. Cells assigned multiple organoid BCs were excluded (<5%), except for those with correlation coefficients > 0.1, which were considered for merging (e.g., organoid BC pairs 1 & 21, 3 & 24, 5 & 22, and 20 & 26 were merged). These likely correspond to cells with more than one gRNA integrant. Cells were then grouped based on their assigned unique organoid BCs or merged organoid BCs, resulting in a total of 20 clonotypes, after excluding clonotypes with fewer than 50 cells (organoid BCs 23 & 25).

### Data processing for the 9TF-monoclonal-arrayed and 8TF-monoclonal-pooled dataset

Cells from each of the 16 10X lanes were processed and underwent quality control separately. The transcriptome data and gRNA data, including UMI counts for individual features (genes or gRNAs) across cells, were imported into Seurat/v5^37^ as two distinct layers. The gRNAs were filtered out if they were expressed in fewer than 1% of cells. The transcriptome data were processed using the same strategy as described above, with adjusted criteria to remove potentially low-quality cells after manually examining the distribution of UMIs and other measurements across cells. Specifically, cells with UMI counts fewer than 1000 (adjusted to 500 or 750 for some lanes), UMI counts exceeding the top 0.5% of total cells, doublet scores calculated by Scrublet over 0.2, MT% over 10% or Ribo% over 50%.

After combining cells from the 16 10X lanes, we applied Seurat/v5^37^ to this dataset, performing conventional single-cell RNA-seq data processing as we did for the ’allTF-mosaic’ dataset. Of note, we identified one cell cluster characterized by low UMI counts and high Ribo%. Moreover, we did not observe highly expressed genes in this cluster corresponding to any known cell types during this developmental stage. Consequently, we removed cells from this cluster and repeated the aforementioned processing steps. Finally, we annotated individual cell clusters using at least two literature-nominated marker genes per cell type label (**Supplementary Table 1**).

### Identifying clonotypes in the 9TF-monoclonal-arrayed and 8TF-monoclonal-pooled datasets

Cells from each experiment were analyzed separately. We first excluded organoid BCs with UMI counts less than 10 or read counts less than 10, and then performed organoid BC corrections with a Hamming distance < 2. We then examined the joint distribution of log2-scaled UMI counts and log2-scaled read counts for individual organoid organoid BCs. Organoid BCs with a log2-scaled read count ≥ 15 were selected (n = 331 for the “arrayed” experiment and n = 293 for the “pooled” experiment), and then assigned to individual cells if the UMI count was ≥ 10 and the proportion of UMI counts within each cell was ≥ 30%.

After excluding cells without any assigned organoid BCs, the UMI count across the organoid BCs for individual cells was normalized by the total count per cell, followed by scaling the normalized counts per organoid BC. Pairwise Pearson correlations were then computed for each pair of organoid BCs. Cells assigned multiple organoid BCs were excluded, except for those with correlation coefficients > 0.1, which were considered for merging. Cells were then grouped based on their assigned unique organoid BC or merged organoid BCs, resulting in a total of 202 clonotypes for 9TF-monoclonal-arrayed and 157 clonotypes for 8TF-monoclonal-pooled, after excluding clonotypes with fewer than 50 cells. Subsequently, dimensionality reduction using PCA was performed on the UMI counts of organoid BCs across cells assigned to individual clonotypes, followed by visualization in a 2D UMAP.

To assign individual clonotypes with a gRNA, we first excluded clonotypes with fewer than 10 cells whose total UMIs of gRNAs were ≥ 10. Subsequently, we calculated the proportions of UMIs across gRNAs for individual cells and assigned a gRNA to a clonotype if its average proportion across cells within that clonotype exceeded 70% (unique assignments). For clonotypes without unique assignments, we also kept their top two most abundant gRNAs (double assignments).

### Hooke analysis

To assess differences in cell abundances between each perturbation and wild type across the dataset, we applied a new computational method, *Hooke*^42^ (https://cole-trapnell-lab.github.io/hooke/), which uses Poisson-lognormal models to account for variability in count data while evaluating the impact of perturbations on relative cell type abundances. We used Hooke to compare abundances across 12 defined cell types between clonotypes assigned to each TF target and NTCs in the arrayed experiment and pooled experiment, respectively. Clonotypes with fewer than 100 cells were excluded from this analysis.

### Differential expressed genes

Differentially expressed genes between Hand1 knockouts and NTC monoclonal EBs were identified using both pseudobulk and single-cell approaches across two experiments (‘9TF-monoclonal-arrayed’ and ‘8TF-monoclonal-pooled’). For the pseudobulk analysis, clonotypes assigned to Hand1 or NTC gRNAs were aggregated and compared using DESeq2^47^, treating individual clonotypes as replicates. For the single-cell analysis, cells were classified as Hand1 or NTC based on gRNA UMI abundance exceeding 90%, and differential expression was assessed using Seurat’s FindMarkers function^37^.

### Processing of ‘9TF-monoclonal-sci’ dataset

Read alignment and gene count matrix generation was performed using the pipeline that we developed for sci-RNA-seq3 (https://github.com/JunyueC/sci-RNA-seq3_pipeline). Briefly, base calls were converted to fastq format using Illumina’s *bcl2fastq*/v2.20 and demultiplexed based on PCR i5 and i7 barcodes using maximum likelihood demultiplexing package *deML* with default settings^68^. Demultiplexed reads were filtered based on the reverse transcription (RT) index and hairpin ligation adaptor index (Levenshtein edit distance (ED) < 2, including insertions and deletions) and adaptor-clipped using *trim_galore*/v0.6.5 (https://github.com/FelixKrueger/TrimGalore) with default settings. Trimmed reads were mapped to the mouse reference genome (mm10) for mouse embryo nuclei using *STAR*/v2.6.1d with default settings^69^ and gene annotations (GENCODE VM12 for mouse). Uniquely mapping reads were extracted, and duplicates were removed using the unique molecular identifier (UMI) sequence, RT index, ligation index and read 2 end-coordinate (that is, reads with identical UMI, RT index, ligation index and tagmentation site were considered duplicates). Finally, mapped reads were split into constituent cellular indices by further demultiplexing reads using the RT index and ligation index. To generate digital expression matrices, we calculated the number of strand-specific UMIs for each cell mapping to the exonic and intronic regions of each gene with *python*/v2.7.13 *HTseq* package^70^. For multi-mapping reads (*i.e.* those mapping to multiple genes), the reads were assigned to the gene for which the distance between the mapped location and the 3’ end of that gene was smallest, except in cases where the read mapped to within 100 bp of the 3’ end of more than one gene, in which case the read was discarded. For most analyses, we included both expected-strand intronic and exonic UMIs in per-gene single-cell expression matrices. After the single cell gene count matrix was generated, cells with low quality (UMI < 100 or detected genes < 100 or unmatched_rate (proportion of reads not mapping to any exon or intron) ≥ 0.4) were filtered out. Each cell was assigned to its originating sample (arrayed_gravity, arrayed_filter, pooled_gravity, and pooled_filter) on the basis of the reverse transcription barcode.

We performed two steps with the goal of exhaustively detecting and removing potential doublets. Of note, all these analyses were performed separately on data from each experiment. First, we used Scrublet^65^ to detect doublets directly. In this step, we first randomly split the dataset into multiple subsets (six for most of the experiments) in order to reduce the time and memory requirements. We then applied the *scrublet*/v0.1 pipeline to each subset with parameters (min_count = 3, min_cells = 3, vscore_percentile = 85, n_pc = 30, expected_doublet_rate = 0.06, sim_doublet_ratio = 2, n_neighbors = 30, scaling_method = ’log’) for doublet score calculation. Cells with doublet scores over 0.2 were annotated as detected doublets. Second, we performed two rounds of clustering and used the doublet annotations to identify subclusters that are enriched in doublets. The clustering was performed based on *Scanpy*/v.1.6.0^71^. Briefly, gene counts mapping to sex chromosomes were removed, and genes with zero counts were filtered out. Each cell was normalized by the total UMI count per cell, and the top 3,000 genes with the highest variance were selected, followed by renormalizing the gene expression matrix. The data was log-transformed after adding a pseudocount, and scaled to unit variance and zero mean. The dimensionality of the data was reduced by PCA (30 components), followed by Louvain clustering with default parameters (resolution = 1). For the Louvain clustering, we first computed a neighborhood graph using a local neighborhood number of 50 using *scanpy.pp.neighbors*. We then clustered the cells into sub-groups using the Louvain algorithm implemented by the *scanpy.tl.louvain* function. For each cell cluster, we applied the same strategies to identify subclusters, except that we set resolution = 3 for Louvain clustering. Subclusters with a detected doublet ratio (by *Scrublet*) over 15% were annotated as doublet-derived subclusters. We then removed cells which are either labeled as doublets by *Scrublet* or that were included in doublet-derived subclusters. Altogether, 7.5% of cells identified as potential doublets were removed through this procedure.

After removing the potential doublets detected by the above two steps, we further filtered out the potential low-quality cells with the doublet score (by *Scrublet*) greater than 0.15, the percentage of reads mapping to ribosomal chromosomes (Ribo%) > 10, or the percentage of reads mapping to mitochondrial chromosomes (Mito%) > 10. Subsequently, we removed the bottom and top 2.5% of cells based on their UMI counts. The final dataset comprises 46,754 cells, with a median UMI count of 505 and a median gene detection count of 427 per cell.

We then applied Seurat/v5^46^ to this final dataset, performing conventional single-cell RNA-seq data processing: 1) retaining protein-coding genes, lincRNA, and pseudogenes for each cell and removing gene counts mapping to sex chromosomes; 2) normalizing the UMI counts by the total count per cell followed by log-transformation; 3) selecting the 2,500 most highly variable genes and scaling the expression of each to zero mean and unit variance; 4) applying PCA and then using the top 30 PCs to calculate a neighborhood graph, followed by louvain clustering (resolution = 1); 4) performing UMAP visualization in 2D space (min.dist = 0.3). For cell clustering, we manually adjusted the resolution parameter towards modest overclustering, and then manually merged adjacent clusters if they had a limited number of differentially expressed genes (DEGs) relative to one another or if they both highly expressed the same literature-nominated marker genes. Subsequently, we annotated individual cell clusters using at least two literature-nominated marker genes per cell type label (**Supplementary Table 1**).

Unlike previous 10X-based methods, we did not directly capture gRNAs. Instead, we sequenced the organoid barcodes and assigned gRNAs to each monoclonal EB using a lookup table containing 30 predefined 8-bp sections of organoid barcodes linked to gRNAs, using only perfect matches. Sequencing data of organoid barcodes were processed as previously described (see Methods section ‘Identifying clonotypes in the 9TF-monoclonal-arrayed and 8TF-monoclonal-pooled datasets’ above). Briefly, organoid BCs with a log2-scaled read count ≥ 10 were selected (n = 350 for the “arrayed” experiment and n = 388 for the “pooled” experiment), and then assigned to individual cells if the UMI count was ≥ 10 and the proportion of UMI counts within each cell was ≥ 30%. After excluding cells without any assigned organoid BCs, the UMI count across the organoid BCs for individual cells was normalized by the total count per cell, followed by scaling the normalized counts per organoid BC. Pairwise Pearson correlations were then computed for each pair of organoid BCs. Cells assigned multiple organoid BCs were excluded, except for those with correlation coefficients > 0.1, which were considered for merging. Cells were then grouped based on their assigned unique organoid BC or merged organoid BCs. In the “arrayed” experiment, 4,315 cells (17.3%) were assigned to 49 clonotypes with at least 50 cells, averaging 88 ± 58 cells and ranging from 50 to 302 cells per clonotype. In the “pooled” experiment, 3,615 cells (16.5%) were assigned to 42 clonotypes with at least 50 cells, averaging 86 ± 41 cells and ranging from 50 to 215 cells per clonotype.

## Supplementary Tables

**Supplementary Table 1.** Marker genes used for mEB cell type annotation.

**Supplementary Table 2.** References highlighting the essential roles of 31 targets in stem cell and EB biology.

**Supplementary Table 3.** The gRNA library targeting genes or NTCs in each screening experiment.

**Supplementary Table 4.** Comparison of log2-scaled fold changes in gRNA frequencies between mESCs and the original plasmid library for CRISPR-cut and CRISPRi.

**Supplementary Table 5.** Comparison of gRNA frequencies between day 21 EBs and the original plasmid library.

**Supplementary Table 6.** Differentially expressed genes were identified by comparing cells with a targeted TF to NTCs in the lateral plate mesoderm.

**Supplementary Table 7.** Comparing cell abundances across 12 identified cell types between clonotypes assigned to each TF target and the NTC was performed using Hooke analysis.

**Supplementary Table 8.** Differentially expressed genes were identified using DESeq2 by comparing pseudobulk profiles of monoclonal EBs assigned to Hand1 and NTC, reporting only results with adjusted p-values <0.05.

## SUPPLEMENTARY FIGURES

**Supplementary Figure 1.**
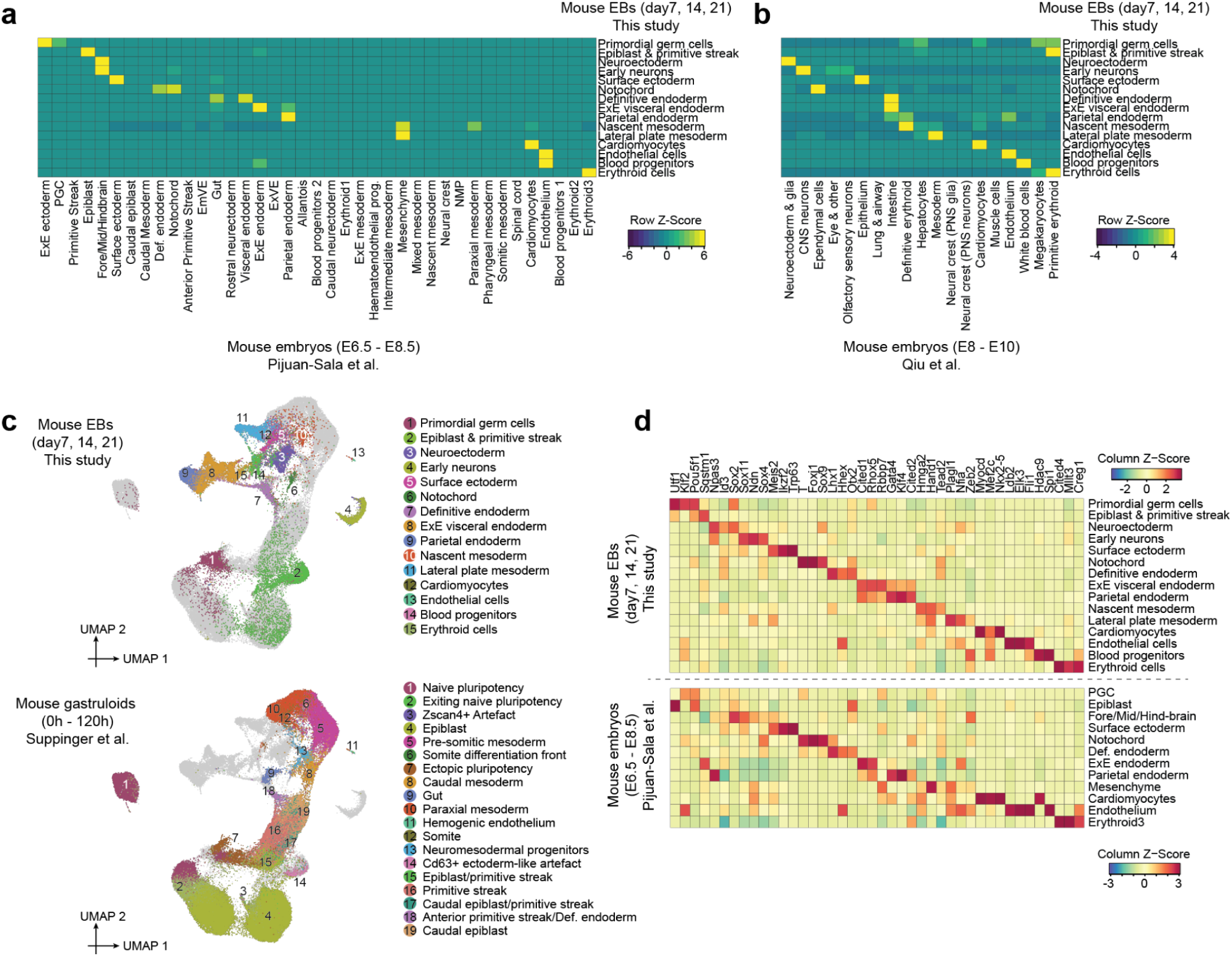
Integrating and co-embedding cells from mouse EBs, embryos, and gastruloids. **a,** Correlated developmental major cell clusters between mouse embryos during gastrulation^23^ (column) and mouse EBs (row) based on non-negative least-squares (NNLS) regression. Heatmap shows the combined regression coefficients (row-scaled) of pairwise cell types between the two datasets. **b,** Correlated developmental major cell clusters between mouse embryos during early somitogenesis^24^ (column) and mouse EBs (row) based on non-negative least-squares (NNLS) regression. Heatmap shows the combined regression coefficients (row-scaled) of pairwise cell types between the two datasets. **c,** UMAP visualization of co-embedded cells from mouse EBs and gastruloids^25^ after batch correction of scRNA-seq data. The same UMAP is shown twice, with colors highlighting cells from either mouse EBs (top) or gastruloids (bottom). **d,** Expression profiles of the top 3 TF markers of the 15 clusters shown in **Fig. 1a**, as identified using the FindAllMarkers function of Seurat/v3^66^, within mouse EBs (top) or mouse embryos during gastrulation^23^ (bottom). Each heatmap illustrates the mean gene expression values within each cluster, calculated from original UMI counts normalized to total UMIs per cell, followed by natural-log transformation. For each cell cluster from the EB dataset, the most similar cell type from mouse embryos was manually selected based on top marker genes, complemented by cell-type correlation analysis, as shown in panel **a**. Overall, we observe that the cell type-specific expression patterns of many of these TFs are shared between related cell types in mouse EBs and gastrulating mouse embryos.

**Supplementary Figure 2.**
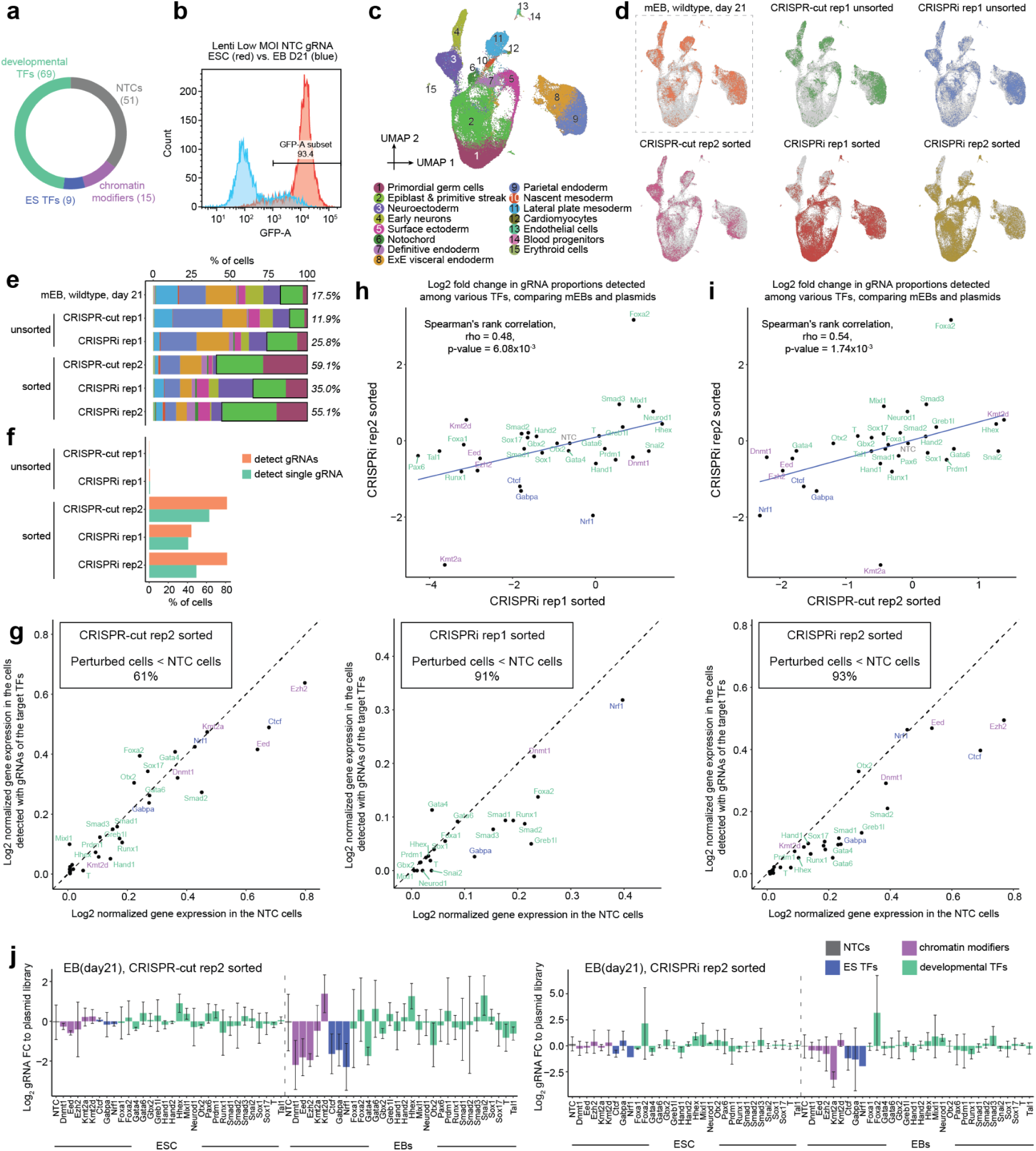
Quality assessment of the CRISPRcut and CRISPRi systems in mouse EBs (‘pilot-mosaic’ experiment). **a,** Pie-chart of pilot library composition. In the pilot screen, a total of 144 gRNAs were designed. Among these, 51 gRNAs corresponded to non-targeting controls (NTC), while the remaining 93 gRNAs were specific to 31 target genes, with 3 gRNAs allocated per target. Target genes were selected to include 3 TFs thought to be essential in mESCs, 5 chromatin modifiers thought to be non-essential in mESCs but required for EB differentiation and 23 TFs with reported roles in germ layer formation. **b,** Exemplary FACS analysis of mESCs (here: CRISPRcut mESCs) transduced with a lentiviral CROPseq library (here: NTC gRNAs), and of 21 day-old EBs generated from these same mESCs, showing loss of GFP expression over the course of differentiation (here: 93% → 21%). Similar ratios were observed for all libraries and cell lines. **c,** 2D UMAP visualization of scRNA-seq profiles from 66,213 co-embedded cells derived from wildtype or perturbed mouse EBs at day 21, following batch correction. These perturbed samples originate from four pilot experiments conducted on mouse EBs at day 21, utilizing either CRISPRcut or CRISPRi techniques, with sorting for GFP-positive cells or without and with two different transduction rates: CRISPRcut rep1 sorted and unsorted (1% GFP positive; not enough cells recovered for sorted population), CRISPRcut rep2 sorted (<20% GFP positive), CRISPRi rep1 sorted and unsorted (1% GFP positive), CRISPRi rep2 sorted (<20% GFP positive). **d,** The same UMAP as in panel **c** is shown multiple times, with colors highlighting cells derived from either wildtype EBs or perturbed EBs from each pilot experiment. **e,** Cell cluster composition of mouse EBs, either from wildtype EBs or perturbed EBs from each pilot experiment. The percentage of less differentiated cells (primordial germ cells, and epiblast & primitive streak) is shown on the right side of each bar. **f,** The percentage of cells with at least one gRNA captured, as well as those with a unique gRNA captured, is reported for each experiment. **g,** Log2-normalized gene expression levels of a particular gene were compared between cells with NTC gRNAs and cells with gRNAs specifically targeting that gene, in three pilot experiments. The percentage of genes with lower expression in perturbed cells compared to NTC cells (below the y = x dotted line) is highlighted in each panel. **h,** Comparison of the log2-fold-change in the proportion of gRNAs targeting a specific gene or NTC between mouse EBs and plasmid library, across two independent CRISPRi pilot experiments. **i,** Comparison of the log2-fold-change in the proportion of gRNAs targeting a specific gene or NTC between mouse EBs and plasmid library, across CRISPRcut and CRISPRi pilot experiments. **j,** Changes in gRNA frequency for different target genes. The log2-fold-change in the proportion of gRNAs was compared between mESCs and plasmid library, or between EBs and plasmid library, in both the CRISPRcut pilot experiment (left) and the CRISPRi pilot experiment (right). Bars represent the average log2-fold-change of gRNAs targeting each gene, with error bars indicating the mean ± standard deviation. In panels **g-j**, target genes are categorized into chromatin modifiers, ES TFs, or developmental TFs.

**Supplementary Figure 3.**
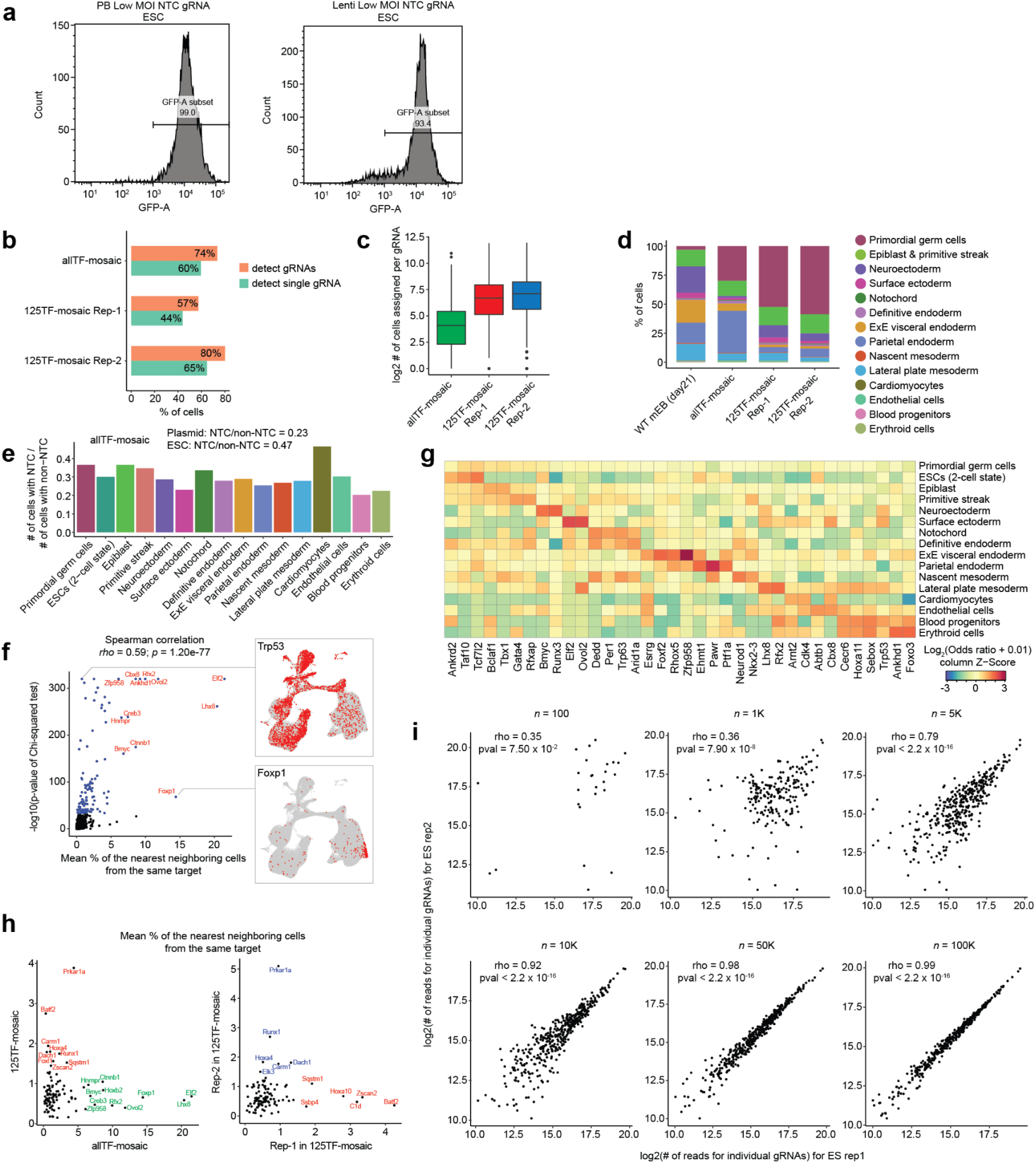
Quality-control of large-scale TF screens in mosaic EBs. **a,** FACS analysis showing high rate of GFP-positive (*i.e.* inferred to be gRNA-expressing) cells after applying puro selection to a starting cell population with <10% GFP-positive cells post-transduction. Due to the combination of low transduction rate and drug selection, the resulting cell population is expected to consist almost entirely of cells expressing exactly one gRNA. **b,** The percentage of cells with at least one gRNA captured, as well as those with a unique gRNA captured, is reported for each of three TF screen datasets. **c,** The number of cells assigned per gRNA were reported for each of the three TF screen datasets (*n* = 4,931 gRNAs for ‘allTF-mosaic’ experiment, and *n* = 433 and 435 for replicates 1 and 2, respectively, of ‘125TF-mosaic’ experiment). Boxplots represent IQR (25th, 50th, 75th percentile) with whiskers representing 1.5× IQR. **d,** Composition of mouse EBs, either from wildtype EBs (21 days) or perturbed EBs from each of the three TF screen datasets, binned by cell cluster. Early neurons, floor plate, and eye field were merged into ’neuroectoderm’, while epiblast, primitive streak, and ESCs (2-cell state) were merged into ’epiblast & primitive streak’ to align with classifications across the different datasets. **e,** Ratio of cells assigned to NTC gRNA versus non-NTC gRNA for each cell type (‘allTF-mosaic’ experiment). **f,** From the ‘allTF-mosaic’ dataset, TF targets with gRNAs detected in fewer than 50 cells were excluded, and cells with each TF target were randomly downsampled to 200 cells. Individual cells were searched for 20-nearest neighbors in PCA space (*n* = 30 dimensions). For each TF target, the average proportion of nearest neighbor cells sharing the same TF target was calculated. These average proportions (x-axis) were then compared with the p-values obtained from chi-squared tests (y-axis) for the same TF targets. Significant TF targets identified by the chi-squared tests are highlighted in blue. Selected TF targets with high values on both axes are labeled. The same UMAP as in **Fig. 2a** is shown twice on the right, with colors highlighting cells where *Trp53* or *Foxp1* transcripts were detected. **g,** From the ‘allTF-mosaic’ dataset, TF targets with gRNAs detected in fewer than 50 cells were filtered out. For each remaining TF target and each of the 16 cell clusters, we performed Fisher’s exact test on the frequency of cells within or outside the cell cluster between cells with the TF target and all cells with NTC gRNAs. The log_2_(odds ratio + 0.01) of the top three TF targets (ranked by p-value) for each cell cluster is presented in the heatmap. **h,** Comparison between experiments of proportion of cells’ nearest neighbors in PCA space sharing the same target TF. For each dataset and for each TF target, the average proportion of nearest neighbor cells within the 20-nearest neighbors in PCA space that shared the same TF target was calculated. Left: comparison of ‘allTF-mosaic’ vs. combined replicates of ‘125TF-mosaic’ experiments. Right: comparison of two replicates of ‘125TF-mosaic’ experiment. Selected TF targets with high proportions in either experiment being compared are labeled. **i,** Standard curve assessing our ability to quantify gRNA read frequency from genomic DNA for different starting cell numbers. Two replicates of variable numbers of mESCs (100, 1K, 5K, 10K, 50K, 100K) containing the 125 TF library (456 distinct gRNAs, with mostly 1 gRNA/cell) were sampled, their genomic DNA was isolated, the gRNA amplified and sequenced. This experiment verified our ability to reproducibly quantify gRNA frequency from as few as 10,000 cells (R=0.92). This serves as a control for quantifying gRNA frequencies from genomic DNA isolated from EB cells containing the same library (each EB contains at least 10K cells, but typically around 30K).

**Supplementary Figure 4.**
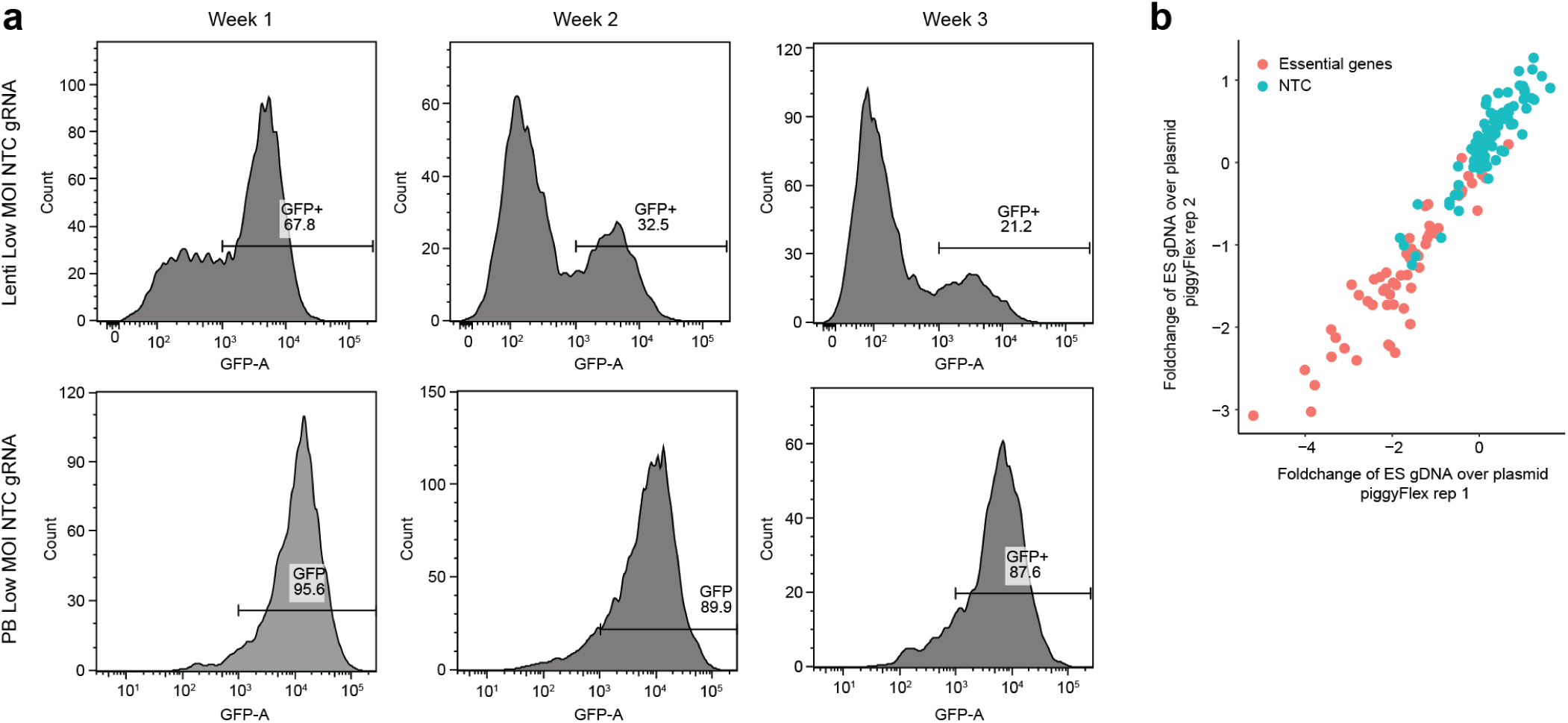
Validation of stable expression during differentiation and functionality of gRNAs with the integrated piggyFlex construct. **a,** FACS analysis plots showing different rates of loss of GFP expression over the course of one, two or three weeks of EB differentiation, for lentiviral CROP-seq vectors (top) and transposon-based piggyFlex (bottom). **b,** Reproducible activity of piggyFlex construct in CRISPRcut mESCs. Fourteen days after transfecting cells with a piggyFlex library of gRNAs containing NTCs and gRNAs targeting essential genes, genomic DNA was isolated and the contained gRNAs amplified by PCR from both the genomic DNA and the plasmid library. Reproducible depletion of gRNAs targeting essential genes (red) compared to NTCs (green) in genomic DNA from two independently transfected replicates implies the CRISPRcut-piggyFlex system is functional.

**Supplementary Figure 5.**
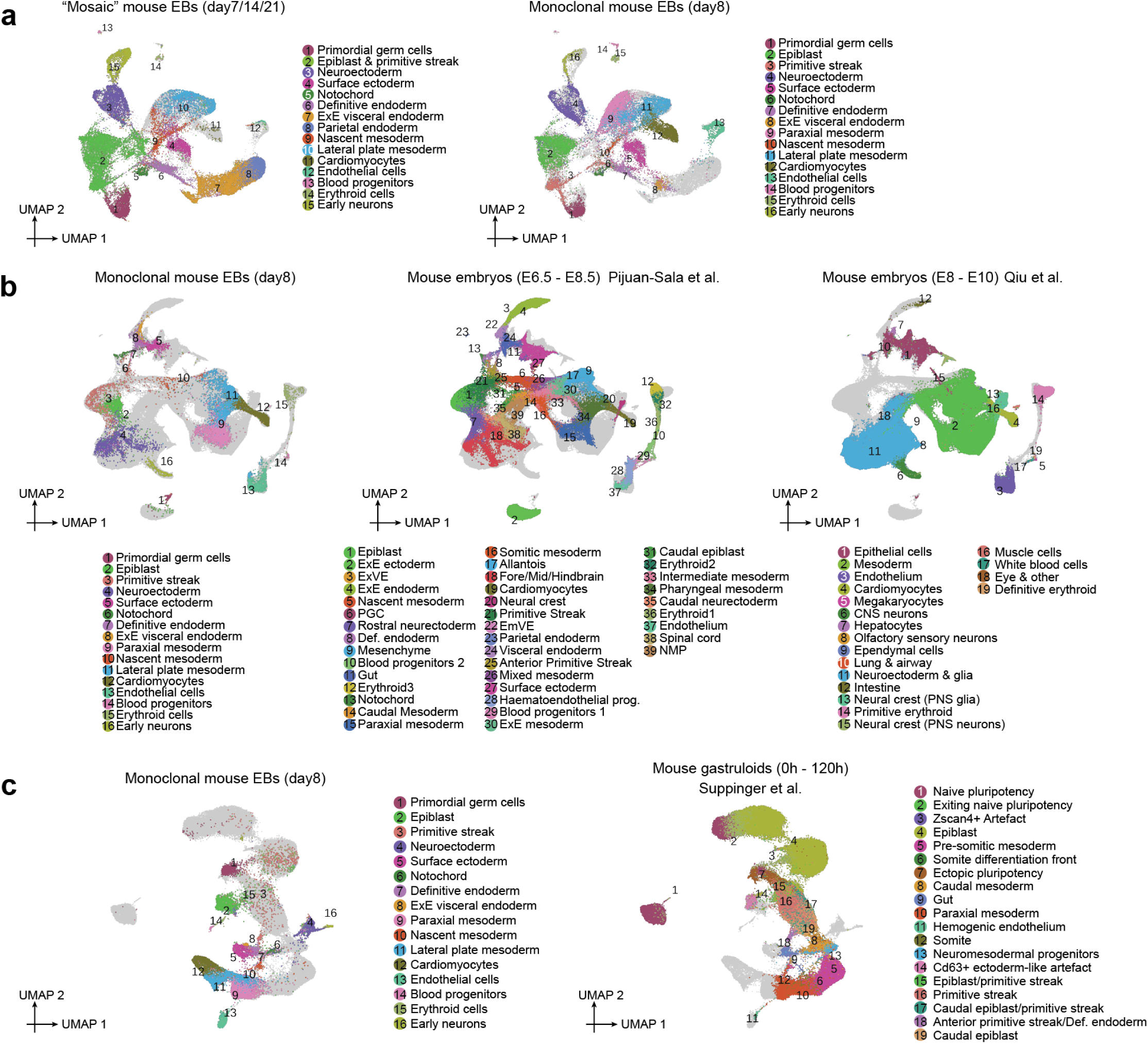
Integrating and co-embedding cells from monoclonal mouse EBs, embryos, and gastruloids. **a,** UMAP visualization of co-embedded cells from ’mosaic’ mouse EBs and monoclonal mouse EBs after batch correction of scRNA-seq data. The same UMAP is shown twice, with colors highlighting cells from either ’mosaic’ mouse EBs (left), or monoclonal mouse EBs (right). **b,** UMAP visualization of co-embedded cells from monoclonal mouse EBs and real embryos at various developmental stages after batch correction of scRNA-seq data. The same UMAP is shown three times, with colors highlighting cells from either mouse EBs (left), embryos during gastrulation^23^ (middle), or embryos during early somitogenesis^24^ (right). **c,** UMAP visualization of co-embedded cells from monoclonal mouse EBs and gastruloids after batch correction of scRNA-seq data. The same UMAP is shown twice, with colors highlighting cells from either mouse EBs (left) or gastruloids^25^ (right).

## REFERENCES

1. Bissiere, S., Gasnier, M., Alvarez, Y. D. & Plachta, N. Cell Fate Decisions During Preimplantation Mammalian Development. Curr. Top. Dev. Biol. 128, 37–58 (2018).

2. Spitz, F. & Furlong, E. E. M. Transcription factors: from enhancer binding to developmental control. Nat. Rev. Genet. 13, 613–626 (2012).

3. Baillie-Benson, P., Moris, N. & Martinez Arias, A. Pluripotent stem cell models of early mammalian development. Curr. Opin. Cell Biol. 66, 89–96 (2020).

4. Chen, B., Du, C., Wang, M., Guo, J. & Liu, X. Organoids as preclinical models of human disease: progress and applications. Med. Rev. 4, 129–153 (2024).

5. Clevers, H. Modeling development and disease with organoids. Cell 165, 1586–1597 (2016).

6. Sato, T. et al. Single Lgr5 stem cells build crypt-villus structures in vitro without a mesenchymal niche. Nature 459, 262–265 (2009).

7. Evans, M. J. & Kaufman, M. H. Establishment in culture of pluripotential cells from mouse embryos. Nature 292, 154–156 (1981).

8. Lewis-Israeli, Y. R. et al. Self-assembling human heart organoids for the modeling of cardiac development and congenital heart disease. Nat. Commun. 12, 5142 (2021).

9. Wagstaff, E. L., Ten Asbroek, A. L. M. A., Ten Brink, J. B., Jansonius, N. M. & Bergen, A. A. B. An alternative approach to produce versatile retinal organoids with accelerated ganglion cell development. Sci. Rep. 11, 1101 (2021).

10. Lancaster, M. A. et al. Cerebral organoids model human brain development and microcephaly. Nature 501, 373–379 (2013).

11. Pettinato, G. et al. Generation of fully functional hepatocyte-like organoids from human induced pluripotent stem cells mixed with Endothelial Cells. Sci. Rep. 9, 8920 (2019).

12. Li, C. et al. Single-cell brain organoid screening identifies developmental defects in autism. Nature 621, 373–380 (2023).

13. Fleck, J. S. et al. Inferring and perturbing cell fate regulomes in human brain organoids. Nature 621, 365–372 (2023).

14. Wahle, P. et al. Multimodal spatiotemporal phenotyping of human retinal organoid development. Nat. Biotechnol. 41, 1765–1775 (2023).

15. Liang, J. et al. In-organoid single-cell CRISPR screening reveals determinants of hepatocyte differentiation and maturation. Genome Biol. 24, 251 (2023).

16. Sheridan, S. D., Surampudi, V. & Rao, R. R. Analysis of embryoid bodies derived from human induced pluripotent stem cells as a means to assess pluripotency. Stem Cells Int. 2012, 738910 (2012).

17. Rhodes, K. et al. Human embryoid bodies as a novel system for genomic studies of functionally diverse cell types. Elife 11, (2022).

18. Kim, I. S. et al. Parallel Single-Cell RNA-Seq and Genetic Recording Reveals Lineage Decisions in Developing Embryoid Bodies. Cell Rep. 33, 108222 (2020).

19. Argelaguet, R. et al. Multi-omics profiling of mouse gastrulation at single-cell resolution. Nature 576, 487–491 (2019).

20. Spangler, A., Su, E. Y., Craft, A. M. & Cahan, P. A single cell transcriptional portrait of embryoid body differentiation and comparison to progenitors of the developing embryo. Stem Cell Res. 31, 201–215 (2018).

21. Juran, C. M., Zvirblyte, J. & Almeida, E. A. C. Differential Single Cell Responses of Embryonic Stem Cells Versus Embryoid Bodies to Gravity Mechanostimulation. Stem Cells Dev. 31, 346–356 (2022).

22. Isakova, A., Neff, N. & Quake, S. R. Single-cell quantification of a broad RNA spectrum reveals unique noncoding patterns associated with cell types and states. Proc. Natl. Acad. Sci. U. S. A. 118, (2021).

23. Pijuan-Sala, B. et al. A single-cell molecular map of mouse gastrulation and early organogenesis. Nature 566, 490–495 (2019).

24. Qiu, C. et al. A single-cell time-lapse of mouse prenatal development from gastrula to birth. Nature 626, 1084–1093 (2024).

25. Suppinger, S. et al. Multimodal characterization of murine gastruloid development. Cell Stem Cell 30, 867–884.e11 (2023).

26. Stuart, T. et al. Comprehensive Integration of Single-Cell Data. Cell 177, 1888–1902.e21 (2019).

27. Qi, L. S. et al. Repurposing CRISPR as an RNA-guided platform for sequence-specific control of gene expression. Cell 152, 1173–1183 (2013).

28. Datlinger, P. et al. Pooled CRISPR screening with single-cell transcriptome readout. Nat. Methods 14, 297–301 (2017).

29. Hong, S. et al. Functional analysis of various promoters in lentiviral vectors at different stages of in vitro differentiation of mouse embryonic stem cells. Mol. Ther. 15, 1630–1639 (2007).

30. Cabrera, A. et al. The sound of silence: Transgene silencing in mammalian cell engineering. Cell Syst 13, 950–973 (2022).

31. Mandegar, M. A. et al. CRISPR interference efficiently induces specific and reversible gene silencing in human iPSCs. Cell Stem Cell 18, 541–553 (2016).

32. Kanamori, M. et al. A genome-wide and nonredundant mouse transcription factor database. Biochem. Biophys. Res. Commun. 322, 787–793 (2004).

33. Huang, X. et al. Single-cell, whole-embryo phenotyping of mammalian developmental disorders. Nature 623, 772–781 (2023).

34. Wu, Q. et al. CARM1 is required in embryonic stem cells to maintain pluripotency and resist differentiation. Stem Cells 27, 2637–2645 (2009).

35. Carpenedo, R. L., Sargent, C. Y. & McDevitt, T. C. Rotary suspension culture enhances the efficiency, yield, and homogeneity of embryoid body differentiation. Stem Cells 25, 2224–2234 (2007).

36. Messana, J. M., Hwang, N. S., Coburn, J., Elisseeff, J. H. & Zhang, Z. Size of the embryoid body influences chondrogenesis of mouse embryonic stem cells. J. Tissue Eng. Regen. Med. 2, 499–506 (2008).

37. Hao, Y. et al. Dictionary learning for integrative, multimodal and scalable single-cell analysis. Nat. Biotechnol. (2023) doi:10.1038/s41587-023-01767-y.

38. Papalexi, E. et al. Characterizing the molecular regulation of inhibitory immune checkpoints with multimodal single-cell screens. Nat. Genet. 53, 322–331 (2021).

39. Herbst, F. et al. Extensive methylation of promoter sequences silences lentiviral transgene expression during stem cell differentiation in vivo. Mol. Ther. 20, 1014–1021 (2012).

40. Chardon, F. M. et al. Multiplex, single-cell CRISPRa screening for cell type specific regulatory elements. Nat. Commun. 15, 8209 (2024).

41. Replogle, J. M. et al. Combinatorial single-cell CRISPR screens by direct guide RNA capture and targeted sequencing. Nat. Biotechnol. 38, 954–961 (2020).

42. Duran, M. et al. A statistical framework for inferring genetic requirements from embryo-scale single-cell sequencing experiments. bioRxivorg 2025.04.03.646654 (2025).

43. Maska, E. L. et al. A Tlx2-Cre mouse line uncovers essential roles for hand1 in extraembryonic and lateral mesoderm. Genesis 48, 479–484 (2010).

44. Lynch, A. T. et al. HAND1 level controls the specification of multipotent cardiac and extraembryonic progenitors from human pluripotent stem cells. EMBO J. (2025) doi:10.1038/s44318-025-00409-0.

45. Cao, J. et al. The single-cell transcriptional landscape of mammalian organogenesis. Nature 566, 496–502 (2019).

46. Martin, B. K. et al. Optimized single-nucleus transcriptional profiling by combinatorial indexing. Nat. Protoc. (2022) doi:10.1038/s41596-022-00752-0.

47. Love, M. I., Huber, W. & Anders, S. Moderated estimation of fold change and dispersion for RNA-seq data with DESeq2. bioRxiv (2014) doi:10.1101/002832.

48. Mossine, V. V., Waters, J. K., Hannink, M. & Mawhinney, T. P. piggyBac transposon plus insulators overcome epigenetic silencing to provide for stable signaling pathway reporter cell lines. PLoS One 8, e85494 (2013).

49. Park, T. S. & Han, J. Y. piggyBac transposition into primordial germ cells is an efficient tool for transgenesis in chickens. Proc. Natl. Acad. Sci. U. S. A. 109, 9337–9341 (2012).

50. Coucouvanis, E. & Martin, G. R. Signals for death and survival: a two-step mechanism for cavitation in the vertebrate embryo. Cell 83, 279–287 (1995).

51. Pettinato, G., Wen, X. & Zhang, N. Formation of well-defined embryoid bodies from dissociated human induced pluripotent stem cells using microfabricated cell-repellent microwell arrays. Sci. Rep. 4, 7402 (2014).

52. Ringel, T. et al. Genome-scale CRISPR screening in human intestinal organoids identifies drivers of TGF-β resistance. Cell Stem Cell 26, 431–440.e8 (2020).

53. Rosen, L. U., et al. Inter-gastruloid heterogeneity revealed by single cell transcriptomics time course: implications for organoid based perturbation studies. bioRxiv 2022.09.27.509783 (2022) doi:10.1101/2022.09.27.509783.

54. Regalado, S. G. et al. Lineage recording in monoclonal gastruloids reveals heritable modes of early development. bioRxivorg (2025/4).

55. Choi, J. et al. A time-resolved, multi-symbol molecular recorder via sequential genome editing. Nature 608, 98–107 (2022).

56. Antón-Bolaños, N. et al. Brain Chimeroids reveal individual susceptibility to neurotoxic triggers. Nature 631, 142–149 (2024).

57. Niimi, M. et al. Single embryonic stem cell-derived embryoid bodies for gene screening. Biotechniques 38, 349–50, 352 (2005).

58. Kingdom, R. & Wright, C. F. Incomplete penetrance and variable expressivity: From clinical studies to population cohorts. Front. Genet. 13, 920390 (2022).

59. Gilbert, L. A. et al. CRISPR-mediated modular RNA-guided regulation of transcription in eukaryotes. Cell 154, 442–451 (2013).

60. Doench, J. G. et al. Optimized sgRNA design to maximize activity and minimize off-target effects of CRISPR-Cas9. Nat. Biotechnol. 34, 184–191 (2016).

61. Horlbeck, M. A. et al. Compact and highly active next-generation libraries for CRISPR-mediated gene repression and activation. Elife 5, (2016).

62. Hill, A. J. et al. On the design of CRISPR-based single-cell molecular screens. Nat. Methods 15, 271–274 (2018).

63. Gaidatzis, D., Lerch, A., Hahne, F. & Stadler, M. B. QuasR: quantification and annotation of short reads in R. Bioinformatics 31, 1130–1132 (2015).

64. Zheng, G. X. Y. et al. Massively parallel digital transcriptional profiling of single cells. Nat. Commun. 8, 14049 (2017).

65. Wolock, S. L., Lopez, R. & Klein, A. M. Scrublet: Computational Identification of Cell Doublets in Single-Cell Transcriptomic Data. Cell Syst 8, 281–291.e9 (2019).

66. Hafemeister, C. & Satija, R. Normalization and variance stabilization of single-cell RNA-seq data using regularized negative binomial regression. Genome Biol. 20, 296 (2019).

67. Alexa, A. & Rahnenführer, J. Gene set enrichment analysis with topGO. Bioconductor Improv 27, 1–26 (2009).

68. Renaud, G., Stenzel, U., Maricic, T., Wiebe, V. & Kelso, J. deML: robust demultiplexing of Illumina sequences using a likelihood-based approach. Bioinformatics 31, 770–772 (2015).

69. Dobin, A. et al. STAR: ultrafast universal RNA-seq aligner. Bioinformatics 29, 15–21 (2013).

70. Anders, S., Pyl, P. T. & Huber, W. HTSeq--a Python framework to work with high-throughput sequencing data. Bioinformatics 31, 166–169 (2015).

71. Wolf, F. A., Angerer, P. & Theis, F. J. SCANPY: large-scale single-cell gene expression data analysis. Genome Biol. 19, 15 (2018).

72. McFaline-Figueroa, J. L. et al. A pooled single-cell genetic screen identifies regulatory checkpoints in the continuum of the epithelial-to-mesenchymal transition. Nat. Genet. 51, 1389–1398 (2019).

